# Use of the Polo-like kinase 4 (PLK4) inhibitor centrinone to investigate intracellular signaling networks using SILAC-based phosphoproteomics

**DOI:** 10.1101/2020.05.22.110767

**Authors:** Dominic P Byrne, Christopher J Clarke, Philip J Brownridge, Anton Kalyuzhnyy, Simon Perkins, Amy Campbell, David Mason, Andrew R Jones, Patrick A Eyers, Claire E Eyers

## Abstract

Polo-like kinase 4 (PLK4) is the master regulator of centriole duplication in metazoan organisms. Catalytic activity and protein turnover of PLK4 are tightly coupled in human cells, since changes in PLK4 concentration and catalysis have profound effects on centriole duplication and supernumerary centrosomes, which are associated with aneuploidy and cancer. Recently, PLK4 has been targeted with a variety of small molecule kinase inhibitors exemplified by centrinone, which rapidly induces inhibitory effects on PLK4 and leads to on-target centrosome depletion. Despite this, relatively few PLK4 substrates have been identified unequivocally in human cells, and PLK4 signaling outside centriolar networks remains poorly characterised. We report an unbiased mass spectrometry (MS)-based quantitative analysis of cellular protein phosphorylation in stable PLK4-expressing U2OS human cells exposed to centrinone. PLK4 phosphorylation was itself sensitive to brief exposure to the compound, resulting in PLK4 stabilization. Analysing asynchronous cell populations, we report hundreds of centrinone-regulated cellular phosphoproteins, including centrosomal and cell cycle proteins and a variety of likely ‘non-canonical’ substrates. Surprisingly, sequence interrogation of ~300 significantly down-regulated phosphoproteins reveals an extensive network of centrinone-sensitive [Ser/Thr]Pro phosphorylation sequence motifs, which based on our analysis might be either direct or indirect targets of PLK4. In addition, we confirm that NMYC and PTPN12 are PLK4 substrates, both *in vitro* and in human cells. Our findings suggest that PLK4 catalytic output directly controls the phosphorylation of a diverse set of cellular proteins, including Pro-directed targets that are likely to be important in PLK4-mediated cell signaling.

## INTRODUCTION

Polo-like kinases (PLKs) are key cell-cycle Ser/Thr kinases conserved in metazoan organisms. Four kinase-domain-containing PLK family members are found in most kinomes, and each PLK polypeptide is thought to serve specific functions in cells. Polo-like kinase 4 (PLK4) is the central regulator of centriole assembly [1, 2]. Specifically, PLK4 activity is rate-limiting for centriole duplication, a process that requires hierarchical recruitment of a number of evolutionarily-conserved core proteins: PLK4/SAK/STK18/ZYG-1 itself, SAS-5/Ana2/STIL, SPD-2/DSpd-2/CEP192, Sas6, and Sas4/CPAP to a single assembly site on the mother centriole [3], driven by its role in phosphorylating key components of the centriolar duplication machinery, notably STIL. Overexpression of PLK4 induces centrosome amplification from pre-existing centrioles and drives *de novo* centriole assembly [1–8]. In human cells, PLK4 is recruited to the centriole during G1 phase through interaction with CEP152 and CEP192. At the G1/S transition, PLK4 transforms from a ring-like localization to a single focus on the wall of the parent centriole that marks the site of procentriole formation [9–12]. Binding of PLK4 to the physiological centriolar substrate STIL promotes activation of PLK4, and the subsequent binding and recruitment of SAS6 [13–16].

Distinct from its canonical rate-limiting role in the control of centriolar duplication, non-centriolar PLK4 has also been implicated in actin-dependent cancer cell migration and invasion, cell protrusion, and invasion and metastasis in model cancer xenografts. Mechanistically, PLK4 functionally targets the Arp2/3 complex and a physical and functional interaction between PLK4 and Arp2 drives PLK4-driven cancer cell movement [17–19]. An interaction between STIL, CEP85 and PLK4 is also implicated in cytoskeletal dynamics [20], and the WNT signaling pathway represents another recently described non-canonical PLK4 target [21].

Like many Ser/Thr protein kinases, PLK activity is itself controlled by phosphorylation in the activation segment; for PLK1 this is driven through Aurora A-dependent phosphorylation at Thr210 in the PLK1 T-loop [22, 23]. In contrast, PLK4 autoactivates through template-driven autophosphorylation in its activation segment, where at least six sites of autophosphorylation, notably trans-autophosphorylated Thr170 (Thr172 in flies) [4], are conserved across multiple species [6, 24, 25]. To evaluate potential PLK4 substrates, the active human PLK4 catalytic domain can be conveniently expressed in bacteria, where autoactivation is also mediated by autophosphorylation at multiple activation segment amino acids, including a non-canonical Tyr residue [26, 27]. PLK4 possesses a triple polo box architecture that facilitates oligomerization, centriole and substrate targeting [28], and helps promote *trans*-autophosphorylation of PLK4 at multiple amino acids [29–32]. The polo-box domains (PBDs) of human PLK1-3, and those from budding yeast CDC5 and Plx1 from *Xenopus laevis*, all recognize similar pS/pT-binding motifs in a proline-directed S[pS/pT]P consensus, which allows PLKs to interact with pre-phosphorylated ‘primed’ substrates as part of ordered enrichment mechanisms in cells [33].

Binding of the PLK4 substrate STIL promotes a key auto-phosphorylation event within the 23 amino acid phosphodegron (residues 282-305) [13]. PLK4 ubiquitination then ensues, targeting it for proteosomal-mediated degradation [6, 24, 25]. This process of self-destruction ensures that centriole duplication is limited to only once per cell cycle and provides an additional mechanism for temporal control of phosphorylation of downstream targets, both directly or through priming phosphorylation events [6, 24, 25, 34]. Although additional physiological PLK4 substrates have been identified [6, 14, 35–38], a detailed characterisation of cellular PLK4 substrates phosphorylated during G1/S phase, or indeed across the phases of the cell cycle, is still lacking. Consequently, the contribution of PLK4 to centrosomal and non-centrosomal biology, ciliopathies and congenital abnormalities associated with gene dosage affects, alongside PLK4 kinase-independent functions that stabilize the kinase, remain to be established [6, 39–41].

PLK substrate specificity has been evaluated using both proteins and peptide arrays, and a broad acidic consensus for phosphorylation by PLK1 has been established: [D/E]x[pS/pT]Φ, where x is any amino acid andis an amino acid with a hydrophobic side chain [42, 43]. PLK2 and PLK3 also appears to possess a preference for acidic residues adjacent to the site of phosphorylation [44], with some bias for *N*- and *C*-terminal negative charge in the canonical [D/E]x[pS/pT]xEE motif [45, 46]. In contrast, PLK4 is unusual in the PLK family with respect to both substrate and small molecule specificity, and sequence and substrate similarity is highest between PLK2 and PLK3, followed by PLK1 and then PLK4, which is the least representative of the family [45]. Consistently, a dual *C*-terminal hydrophobic consensus ([pS/pT]ΦΦ) has been ascribed to efficient PLK4 substrate phosphorylation in the literature, based on specificity studies using (PLK1)-based optimised synthetic peptide arrays [47] and a variety of recombinant protein assays [16, 38]. Consistently, many physiological PLK4 substrates conform to this broad hydrophobic consensus *C*-terminal to the site of modification. However, other variants of peptide substrates have also been shown to be phosphorylated by PLK4 in peptide array studies, with bias shown towards those with an Asp or Asn at the −2 position [45], as well as hydrophobic and basic amino acids *C*-terminal to the site of phosphorylation [45–47]. Justifiably, these findings have guided the research community in the search for distinct PLK1, 2, 3 and 4 substrates. Indeed, chemical-genetic screens and phosphospecific antibodies, have led to the identification of a variety of cellular PLK4 substrates, including STIL, CEP192 and CEP131, in vertebrate cells [16, 38]. While these studies also uncovered a number of potential additional PLK4-dependent phosphorylation sites in presumed targets, interpretation of these data were guided by their potential localisation to the centrosome, and/or an involvement in centrosomal biology, usually with the explicit assumption that PLK4 phosphorylates sites in substrates conform to a ([pS/pT]ΦΦ) consensus.

The relatively high specificity of the cell permeable PLK4 inhibitor centrinone [48] has led to its adoption as a tool compound for rapid PLK4 inactivation in living cells. Unlike promiscuous PLK4 inhibitors such as CFI-400945 [49], centrinone was chemically-optimised using the pan-Aurora kinase and PLK4 inhibitor VX-680 (MK-0457) [50], as the template. The addition of a methoxy substituent at the C5 position promotes interaction with Met91, lying 2 amino acids C-terminal from the gatekeeper residue Leu89 in the PLK4 hinge region [51]. This results in a reported 2 orders-of-magnitude increase in inhibition of PLK4 relative to Aurora A. Moreover, incubation of human cells with centrinone provides a useful approach for studying a variety of phenotypic effects [41, 52], with the explicit caveat that, as with essentially all kinase inhibitors, off-target effects are inevitable. Most notably, prolonged centrinone exposure leads to the depletion of centrosomes, causing cell-cycle arrest in G1 via a p53-dependent mechanism, potentially by loss of the interaction between p53 and the ubiquitin E3 ligase MDM2 [48]. Importantly, centrinone exposure of cells containing a drug-resistant PLK4 (in which Gly95, which is critical for drug-binding, is mutated to Leu) provided compelling evidence that the loss of centrosome phenotype is a direct result of PLK4 inhibition, since centrosomes are maintained normally in the presence of centrinone in the drug-resistant cell line [48]. Experimentally, the differing substrate-specificity of Aurora kinases, which are primarily basophilic Ser/Thr kinases that do not tolerate Pro in the +1 position [53, 54], and PLK4, which has no obvious preference for Arg/Lys *N*-terminal to the site of substrate phosphorylation [45, 47], can also simplify the analysis of phosphoproteomics datasets generated with specific small molecules such as centrinone.

In this study, we perform an in-depth quantitative phosphoproteomics analysis to identify, in an unbiased manner, putative direct and downstream targets of PLK4. By exploiting centrinone and a previously described drug-resistant G95R PLK4 allele [55], we identify hundreds of centrinone-regulated phosphorylation sites in human U2OS cells that are sensitive to acute drug exposure. Surprisingly, phosphorylation motif analysis reveals a Pro-directed consensus in the majority of centrinone-sensitive phosphosites, which might therefore be dependent on PLK4 catalytic activity. Consistently, we confirm that three centrinone targets, the tyrosine phosphatase PTPN12, the neuroblastoma-driving transcription factor NMYC and PLK4 itself, are directly phosphorylated by recombinant PLK4 *in vitro*. We conclude that non-canonical Pro-directed phosphorylation sites that lie downstream of centrinone have the potential to be direct, or indirect, PLK4 targets, depending upon the substrate and context.

## MATERIALS AND METHODS

### Small molecules, reagents and Antibodies

Unless otherwise stated, general lab reagents were purchased from Sigma Aldrich and were of the highest quality available. The following antibodies were used: anti-PLK4 antibody clone 6H5 (Millipore, used at 1/100 for western blot and immunofluorescence), anti-Aurora A (Cell Signaling Technologies, 1/5000 for western blot, 1/500 for immunofluorescence), anti-pericentrin (Abcam, 1/500 for immunofluorescence), anti-γ-tubulin (1/500 for immunofluorescence), anti-9E10 anti-myc (Invitrogen, 1/100 for immunofluorescence), anti-FLAG (Sigma, 1/1,000 for western blot), anti-α-tubulin (TAT1) (1/5000 for western blot), anti-GAPDH (1/5000 for western blot). N-terminally FITC-coupled (fluorescent) PLK4, NMYC, PTPN12 and CEP131 substrate peptides were designed for EZ reader assays in-house and purchased from Pepceuticals, Leicester, UK (see Table 1 for parental peptides). Centrinone and VX-680 were obtained as previously described [56].

**Table 1.** PLK4 enzyme assays and substrates. Sequence of recombinant protein kinases and peptide substrates employed for assay of recombinant human PLK4. Sources of enzymes are included. 5FAM=5-carboxyfluorescein.

| Substrate sequence | Parental substrate sequence |
| --- | --- |
| **Human PLK4 (bacteria)** | Amino acids 1-269 |
| **Human PTPN12 (wheat germ cell free)** | Amino acids 1-780 |
| **Human NMYC (wheat germ cell free)** | Amino acids 1-464 |
| **PLK4 CDK7/CAK substrate** | 5FAM-FLAKSFGSPNRAYKK-CONH<sub>2</sub> |
| **PLK4 polybasic ('RK') substrate** | 5FAM-RKKKSFYFKKHHH-CONH<sub>2</sub> |
| **NMYC substrate** | 5FAM-KFELLPTPPLSPSR-CONH<sub>2</sub> |
| **PTPN12 substrate** | 5FAM-TVSLTPSPTTQVET-CONH<sub>2</sub> |
| **CEP131 substrate 1** | 5FAM-NLRRSNSTTQVSQ-CONH<sub>2</sub> |
| **CEP131 substrate 2** | 5FAM-SQPRSGSPRPTEP-CONH<sub>2</sub> |

### Cell culture and generation of stable cell lines

U2OS T-REx parental Flp-In cells were maintained in DMEM supplemented with 10% (v/v) foetal bovine serum, penicillin (100 U/mL), streptomycin (100 U/mL) and L-glutamine (2 mM), 0.1 mg/ml Zeocin and 15 μg/ml blasticidin at 37 °C, 5% CO_2_. Stable cells were generated based on published protocols using the Flp recombinase plasmid pOG44 [2, 7]. Briefly, tetracycline-inducible lines expressing either MYC or FLAG-tagged PLK4 were initially generated by transfection of parental T-REx cells (A kind gift of Dr Gopal Sapkota, University of Dundee). Stable U2OS clonal populations that contained either FLAG-WT PLK4, FLAG G95R PLK4, MYC-WT PLK4, or MYC-G95R PLK4 were established by selection for 2-3 weeks with 15 μg/ml blasticidin and 100 μg/ml hygromycin B, as described for other U2OS models [55, 57–59], and FLAG-PLK4 or MYC-PLK4 expression was induced by the addition of 1 μg/ml tetracycline. Cells were maintained in culture and split at ~80 % confluence, when cells were washed with PBS, trypsinised (0.05 % (v/v) and harvested by centrifugation at 220 x*g*.

### Immunofluorescence Microscopy

U2OS FLAG-WT PLK4, FLAG G95R PLK4, MYC-WT PLK4 or MYC-G95R PLK4 stably transfected cells were split into a 6 well plate containing cover slips. At ~80 % confluence, 1 μg/mL tetracycline was added to induce protein expression for 18 hours. Cells were washed with 2.5 mL PBS and fixed for 20 min with 2.5 mL of 3.7% (v/v) paraformaldehyde. Cells were then washed 3x with 2.5 mL PBS and permeabilised with 0.1 % (v/v) Triton X-100 for 10 min. Cells were washed a further 3x with PBS and blocked with 1% (w/v) BSA in 0.3 M glycine for 60 min, and washed twice with PBS. Primary antibodies, anti-9E10 anti-myc antibody (1/100), anti-γ-tubulin (1/5000) or anti-pericentrin (1/500) in 500 μL of 1% (w/v) BSA were added to each coverslip and incubated overnight at 4 °C in a humidified chamber. Cells were washed 5x for 10 min with PBS, prior to incubation for 1 hr (RT, dark) with secondary antibody (rabbit anti-mouse-Cy3 or goat anti-rabbit-FITC, 1 in 200 dilution). Cells were washed 5x 5 min with PBS and incubated with 1 μg/mL DAPI for 3 min. Coverslips were mounted onto microscope slides and secured with immune-mount and imaged on a Zeiss LSM 880 confocal microscope using an alpha Plan-Apochromat 100x/1.46 DIC grade oil immersion lens. A 405 nm diode, 488 nm Argon and 561 nm diode laser were used to excite DAPI, Alexa 488 and Cy3 respectively with a transmitted light image also captured. A pinhole of ~1 Airy unit was maintained throughout imaging. Spatial sampling was conducted using Nyquist sampling for the chosen magnification with four times line averaging used to reduce noise. For each condition at least one z-stack through the sample was acquired with slices taken at ~1 μm throughout the cell under investigation.

### Flow Cytometry (FACS)

U2OS FLAG-WT and G95R PLK4 cells were split into 10 cm^2^ dishes. At 60 % confluence, cells were incubated with 1 μg/mL tetracycline (18 h) before addition of centrinone (300 nM) or vehicle control for 4 hours. Cells were washed with PBS and released with trypsin (0.05 % (v/v)). Following centrifugation at 220 x*g*, PBS was removed and cells fixed with 70 % (v/v) ethanol, added slowly whilst vortexing to avoid aggregation and stored at −20 °C overnight. Fixed cells were washed with PBS and incubated with 200 μL Guava cell cycle reagent for 30 min (RT, dark). Samples were transferred to a 96-well plate and analysed using a Guava easycyte HT cytometer. ANOVA statistical analysis and Tukey post-hoc testing was performed in R.

### SILAC labelling

U2OS T-REx Flp-in cells stably transfected with FLAG-WT PLK4 or FLAG-G95R PLK4 were grown in DMEM supplemented with 10% (v/v) dialysed foetal bovine serum, penicillin (100 U/mL) and streptomycin (100 U/mL). Once 80 % confluency was reached, cells were split in to DMEM containing ‘heavy’ labelled, ^15^N_2_^13^C_6_-lysine (Lys8) and ^15^N_2_^13^C_6_-arginine (Arg10) for ~seven cell doublings to permit full incorporation of the label. At 80 % confluence, cells were washed with PBS, released with trypsin (0.05 % (v/v)) and centrifuged at 220 x*g*.

### Cell lysis

For immunoblotting, cells were lysed in 1% (v/v) NP-40, 1% (w/v) SDS dissolved in 50 mM Tris-HCl pH 8.0 and 150 mM NaCl, supplemented with Roche protease inhibitor cocktail tablet. The lysate was sonicated briefly and centrifuged at 15,000 x*g* (4 °C) for 20 min. Protein concentration was quantified using the Bradford Assay (BioRad). For co-immunoprecipitation (IP) experiments, cells were lysed in 50 mM Tris-HCl (pH 8.0), 150 mM NaCl, 0.5% NP-40, 1 mM DTT, 2 mM MgCl_2_, and benzonase supplemented with a protease inhibitor (Roche) cocktail tablet. After 30 min incubation on ice, cell lysates were clarified by centrifugation (15,000 x*g*, 4 °C, 20 min). For LC-MS/MS analysis, cells were lysed with an MS compatible buffer (0.25% (v/v) RapiGest SF (Waters) in 50 mM ammonium bicarbonate, supplemented with PhosStop inhibitor (Roche). Lysates were sonicated briefly to shear DNA and centrifuged (15,000 x*g*, 4 °C, 20 min).

### FLAG-PLK4 Immunoprecipitation

Anti-FLAG M2 Affinity Agarose resin (Sigma, ~30 μL) was washed 3x with 50 mM Tris-HCl (pH 8.0) and 150 mM NaCl and then incubated with clarified cell lysate overnight at 4 °C with gentle agitation. Agarose beads were collected by centrifugation at 1,000 x *g* for 1 min and then washed 5x with 50 mM Tris-HCl (pH 8.0) containing 150 mM NaCl. Precipitated (bead-bound) proteins were eluted by incubation with 2x SDS sample loading buffer for 5 min at 98 °C.

### Sample preparation for (phospho)proteome analysis

SILAC-labelled protein lysates (1 mg total protein) were mixed with an equal amount of unlabelled protein lysate prior to sample preparation for LC-MS/MS analysis. Proteins were reduced, alkylated, digested with trypsin and desalted using standard procedures [60]. For high pH reversed-phase (RP) HPLC, tryptic peptides were resuspended in 94.5% (v/v) buffer A (20 mM NH4OH, pH 10), 5.5% (v/v) buffer B (20 mM NH4OH in 90% acetonitrile) and loaded on to an Extend C18 (3.5 μm, 3 mm x 150 mm) Agilent column. Peptides were eluted with an increasing concentration of buffer B: to 30% over 25 min, then 75% B over 12 min, at a flow rate of 0.5 mL/min. Sixty 500 μL fractions were collected, partially dried by vacuum centrifugation and concatenated to twelve pools. Aliquots (5 μl) were removed from each of the 12x 50 μl pools for total proteomic analysis, and the remaining sample (45 μl) was subjected to TiO_2_-based phosphopeptide enrichment as previously described [60].

### LC-MS/MS

Reversed-phase capillary HPLC separations were performed using an UltiMate 3000 nano system (Dionex) coupled in-line with a Thermo Orbitrap Fusion Tribrid mass spectrometer (Thermo Scientific, Bremen, Germany) as described [60]. For proteome analysis, full scan MS1 spectra were acquired in the Orbitrap (120k resolution at *m/z* 200) using a top-speed approach (3 s cycle time) to perform for HCD-mediated fragmentation with a normalized collision energy of 32%. MS2 spectra were acquired in the ion trap. Phosphopeptide analysis was performed [60] using the HCD orbitrap method, where both MS1 and MS2 spectra were acquired in the orbitrap (60k resolution for MS1, 30k resolution for MS2).

### (Phospho)proteomics data processing

Phosphopeptide data were processed using Andromeda with PTM-score implemented within MaxQuant (version 1.6.0.16) [61]. MS/MS spectra were searched against a database containing the human UniProt database (downloaded December 2015; 20,187 sequences). Trypsin was set as the digestion enzyme and two missed cleavages were permitted. Cysteine carbamidomethylation was set as a fixed modification. Variable modifications were set as oxidation (M), phospho (S/T/Y). Default instrument parameters and score thresholds were used: MS1 first search peptide tolerance of 20 ppm and main search tolerance of 4.5 ppm; FTMS MS2 tolerance of 20 ppm; ITMS MS2 tolerance of 0.5 Da. A false discovery rate of 1% for peptide spectrum matches (PSM) and proteins was applied. For processing of SILAC data, Thermo files (raw. Format) were loaded in to MaxQuant (version 1.6.0.16). Total proteomics (protein expression data) and phosphoproteomics experiments were processed separately. The experimental template design separated individual bioreplicates into separate experiments, linking the experiment to the relevant fractions. MaxQuant parameters were as described above, with the following additions: ‘multiplicity’ was set to 2 and Arg10/Lys8 were selected as labels. For proteomics datasets, the variable modifications oxidation (M), acetyl (protein *N*-term) were included. Both ‘requantify’ and ‘match between runs’ were enabled. At least two peptides were required for protein quantification. Post processing was performed using Perseus (version 1.6.0.7). For proteingroups.txt output files, Perseus was used to filter out contaminants, reverse decoy hits and those ‘matched only by site’. Additionally, data were filtered to include proteins identified in ≥3 (out of 4) bioreplicates and Log_2_ transformed. For phosphosites(STY).txt output files, data were filtered as above. In addition, ‘expand site table’ feature was used to separate individual phosphosites, and a phosphosite localisation cut-off of ≥0.75 was applied. Ratios were Log_2_ transformed. Statistical analysis was then performed using the LIMMA package in R with Benjamini-Hochberg multiple corrections to generate adjusted p-values.

### Evolutionary conservation

To calculate evolutionary conservation of the most confidently localised phosphosite per phosphopeptide, the sequences of leading assigned proteins for all phosphopeptides were used as a query in a protein-protein BLAST [62] search (BLAST 2.9.0+ version; 11/03/19 build) against the proteomes of 100 eukaryotic species (50 mammals, 12 birds, 5 fish, 4 reptiles, 2 amphibians, 11 insects, 4 fungi, 7 plants and 5 protists) downloaded from UniProt (November 2019) - see supplementary Table 2 for full proteome descriptions. For each target protein, a top orthologue was extracted from each species (E-value <= 0.00001). The orthologues were aligned with the target protein using MUSCLE [63] (version 3.8.31). From the alignments, percentage conservation was calculated for each Ser, Thr and Tyr residue within the sequence of each target protein out of 100 (all proteomes), and out of the number of aligned orthologues. Additional conservation percentages were calculated taking into account Ser/Thr substitutions in orthologues, whereby an orthologue is included in % conservation calculation if, for example, a Thr from an orthologue is aligned with a target Ser and vice versa. Furthermore, conservation percentages were given for −1 and +1 sites around each Ser, Thr and Tyr in target sequence across aligned orthologues. All conservation data was then cross-referenced with phosphopeptide data to determine the conservation of target sites. The most confident phosphosite per phosphopeptide was also cross-referenced against data from PeptideAtlas (PA) [64] (2020 build), and from PhosphoSitePlus (PSP) [65] (11/03/20 build), both of which had been pre-processed to categorise previously observed phosphorylation site confidence categories, based on the number of observations (“High”: ≥ 5 previous observations, very likely true site; “Medium”: 2-4 previous observations, likely true site; “Low”: 1 previous observation, little support that it is a true site; PA only – “Not phosphorylated”: frequently (>5) observed to be not phosphorylated, never observed as phosphorylated; “Other” – no confident evidence in any category). Observations in PA were counted with a threshold of >0.95 PTM Prophet probability for positive evidence, and ≤ 0.19 for evidence of not being phosphorylated.

**Table 2.**
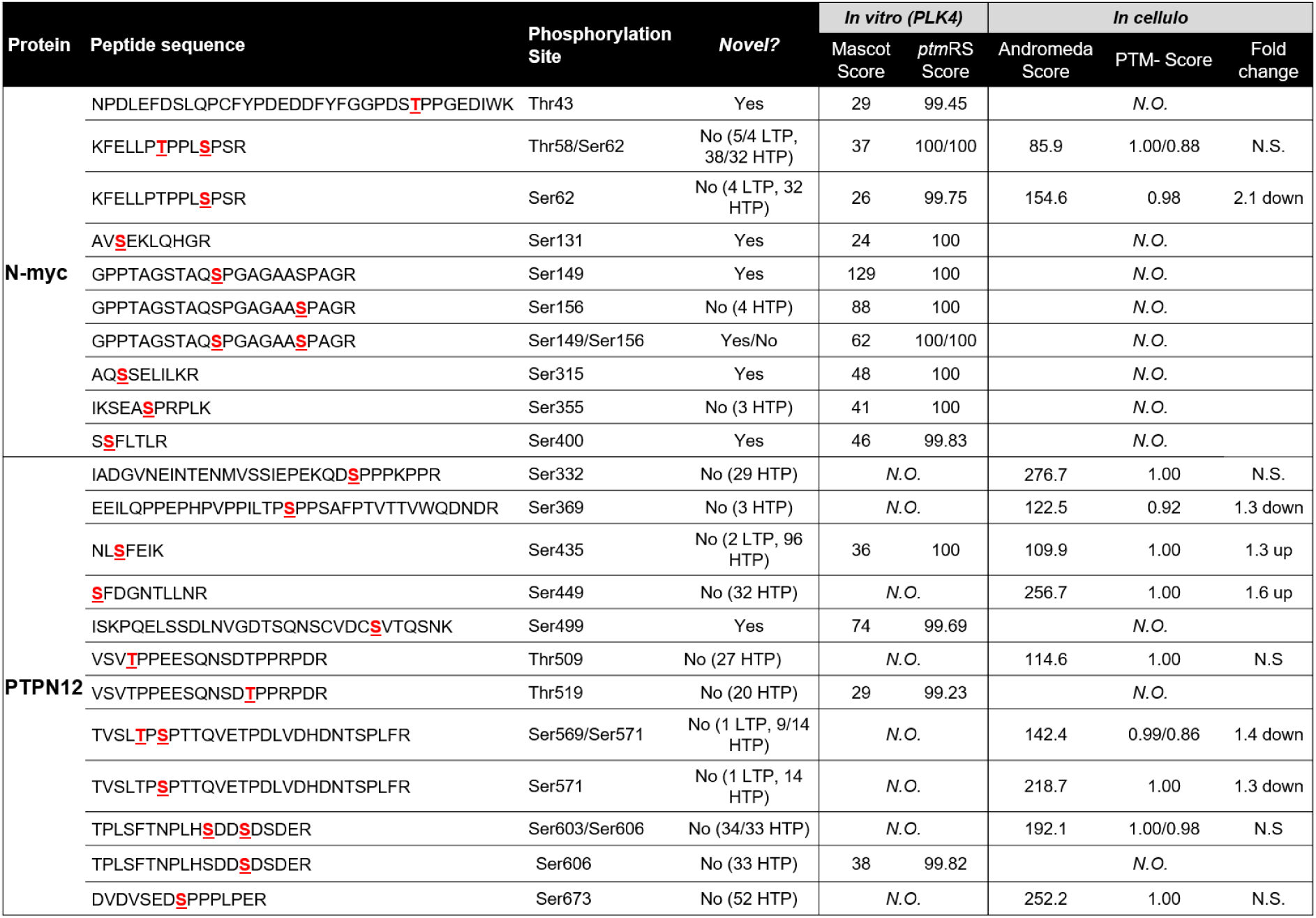
MS-based analysis of NMYC and PTPN12 phosphorylation sites. Phosphopeptides identified from NMYC or PTPN12 by MS/MS following either in vitro phosphorylation by PLK4, or from human cells. The phosphopeptide sequences and the identified site(s) of phosphorylation (underlined within the sequence) are indicated, as well as their Mascot and ptmRS scores (in vitro), or Andromeda and PTM-score (from the cellular analysis), indicating site localisation confidence. Fold-change details the quantitative effect on phosphopeptide levels in response to centrinone. ‘Novelty’ indicates whether these identified phosphosites were previously recorded in PhosphoSitePlus, and whether these records were annotated as either low-throughput (LTP) or high-throughput (HTP) observations. N.O. – not observed; N.S. not significant change in response to centrinone.

### IceLogo analysis

Cellular phosphosites that were modulated after centrinone treatment were aligned to the centre of a sequence window composed of 5 amino acids *N*- and *C*-terminal to the mapped phosphorylation site. Sequence motif logos were generated with IceLogo v1.2 [66] using percentage difference against a background of the precompiled human SwissProt composition and a *p*-value cut-off of 0.05.

### Functional enrichment analysis

Proteins and phosphoproteins that were significantly altered in response to centrinone were subjected to functional enrichment analysis using the DAVID Bioinformatics Resources (v6.8) [67, 68]. “clusterProfiler” R package was used to create dotplots from these datasets, where P‐values, enrichment factor, protein count and functional category could be presented.

### Network analysis

Significantly downregulated proteins from the G95R PLK4 SILAC dataset (adj. *p* value <0.05) were submitted to the STRING database (v. 10.5) to assess protein-protein interactions [69]. A high confidence (score 0.7) filter was applied, and only ‘experiments’ and ‘databases’ as active interaction sources were included.

### Expression and purification of PLK4 catalytic domain

6His-N-terminally tagged human PLK4 catalytic domain (amino acids 1–269) was expressed and purified in bacteria as described previously and affinity purified using Ni-NTA agarose [27]. Proteins were eluted from beads by incubation with buffer containing 0.5 M imidazole. PLK4 amino acid substitutions were introduced using standard PCR-based site-directed mutagenesis protocols and confirmed by sequencing the whole PLK4 1–269 coding region.

### DSF (differential scanning fluorimetry) assays

Thermal shift assays were performed on a StepOnePlus Real-Time PCR machine (Life Technologies) in combination with Sypro-Orange dye (Invitrogen) and a thermal ramping protocol (0.3°C per minute between 25 and 94°C). Recombinant PLK4 proteins were assayed at 5 μM in 50 mM Tris–HCl (pH 7.4) and 100 mM NaCl in the presence or absence of the indicated concentrations of ligand or inhibitor compound [final DMSO concentration 4% (v/v)]. Data were processed using the Boltzmann equation to generate sigmoidal denaturation curves, and average *T*_m_/Δ*T*_m_ values were calculated as described using GraphPad Prism software [70].

### PLK4 phosphorylation assays

All *in vitro* peptide-based enzyme assays were carried out using the Caliper LabChip EZ Reader platform, which monitors phosphorylation-induced changes in the mobility of a fluorescently labelled PLK4 peptide substrate [27, 71, 72]. To assess PLK4 activity, WT or G95R PLK4 (0.5 μg to 10 μg, depending upon the assay) were incubated with 1 mM ATP (to mimic cellular concentration) and 2 μM of the appropriate fluorescent peptide substrate in 25 mM HEPES (pH 7.4), 5 mM MgCl_2_, and 0.001% (v/v) Brij 35. Centrinone-mediated enzyme inhibition was quantified under identical assay conditions using WT or G95R PLK4 (5 μg), in the presence of increasing concentrations of centrinone (10 nM to 100 μM) by monitoring the generation of phosphopeptide during the assay, by real-time peak integration and quantification of relative amounts of peptide and phosphopeptide. Data were normalised with respect to control assays, with pmoles of phosphate incorporated into the peptide generally limited to <20% of maximum, in order to prevent depletion of ATP, end product effects and to ensure assay linearity. Reactions were pre-incubated for 30 min at 37°C prior to addition of ATP. The following peptide substrates (Table 1) were synthesised, lyophilised, and reconstituted in DMSO prior to use at a fixed concentration in PLK4 kinase assays. The human sequence-derived regions for the peptide substrates are: CDK7/CAK (amino acids 158-169) FLAKSFGSPNRAYKK and point mutants FLAKAFGSPNRAYKK, FLAKSFGSPNAAYKK, and FLAKAFGAPNRAYKK, polybasic ‘RK’ substrate RKKKSFYFKKHHH and F6P substitution RKKKSPYFKKHHH, CEP131 (amino acids 72-84) substrate peptide 1, NLRRSNSTTQVSQ, and point mutant NLRRSNATTQVSQ) and CEP131 (amino acids 83-95) substrate peptide 2, SQPRSGSPRPTEP (and point mutant SQPRSGAPRPTEP), NMYC (amino acids 52-65) substrate, KFELLPTPPLSPSR (and point mutants KFELLPAPPLSPSR, KFELLPTPPLAPSR, KFELLPAPPLAPSR and KFELLPTAPLSASR) and PTPN12 (amino acids 565-578) substrate TVSLTPSPTTQVET(andpointmutantsTVSLAPSPTTQVET, TVSLTPAPTTQVET, TVSLAPAPTTQVET, and TVSLTASATTQVET). Where indicated, phosphorylation of peptides by the active Pro-directed kinase GST-CDK2/Cyclin A2 (100 ng purified from *Sf*9 cells, Sigma) was also confirmed in the presence or absence of the ATP competitive inhibitor, Purvalanol A (100 μM). The phosphorylation kinetics of GST-NMYC (full length, amino acids 1-464) or GST-PTPN12 (amino acids 1-780), both expressed in wheatgerm cell-free systems (Abcam), were measuring through PLK4-dependent phosphate incorporation from gamma-^32^P-labelled ATP. WT PLK4 (2 μg) was incubated with 2 μg of either substrate protein and assayed in 50 mM Tris, pH 7.4, 100 mM NaCl, and 1 mM DTT in the presence of 200 μM ATP (containing 2 μCi ^32^P ATP per assay) and 10 mM MgCl_2_ at 30 °C. The reactions were terminated at the indicated time points by denaturation in SDS sample buffer prior to separation by SDS-PAGE and transfer to nitrocellulose membranes. ^32^P-incorporation into PLK4 (autophosphorylation) was detected by autoradiography. Equal loading of appropriate proteins was confirmed by Ponceau S staining and destaining of the membrane. To evaluate site-specific phosphorylation, trypsin proteolysis, TiO_2_-based phosphopeptide enrichment and liquid chromatography–tandem mass spectrometry (LC-MS/MS) analysis was performed using an Orbitrap Fusion mass spectrometer (Thermo Scientific, Bremen) as previously described [60], with enrichment being performed using Titansphere Phos-TiO tips (GL Sciences), as per the manufacturer’s instructions.

## RESULTS

### Inducible stable U2OS cell lines to investigate dynamic PLK4 signaling using the small molecule inhibitor centrinone

To quantify PLK4 signaling pathways in intact cells, we set out to perform a quantitative investigation of the dynamic phosphoproteome of human cells following treatment with the inhibitor centrinone. Importantly, this chemical has been reported to be a specific inhibitor of PLK4, exhibiting high specificity for this Ser/Thr protein kinase compared to the related kinase Aurora A (Fig. 1A). To evaluate potential off-target effects of centrinone inhibition we initially exploited a chemical genetic approach, generating stable isogenic tetracycline-inducible U2OS cell lines expressing either wild-type (WT), or drug-resistant (G95R), N-terminally epitope tagged PLK4, in the presence of the endogenous PLK4 gene. This strategy allowed us to inducibly control PLK4 protein levels for robust detection and quantification of putative PLK4 substrates, including PLK4 itself, which is expressed at very low (sometimes undetectable) levels in the parental cell line. Mutation of the Gly95 residue of PLK4 to Arg results in a catalytically active enzyme that is resistant to concentrations of up to 100 µM centrinone *in vitro*, in the presence of cell-mimicking concentrations (1 mM) of the ATP co-factor (Supplementary Fig. 1A-D). Under these experimental conditions, which are designed to mimic the cellular environment, centrinone possessed an IC_50_ of ~330 nM for WT PLK4 (Supplementary Fig. 1D). Consistently, generation of PLK4 (1-269) containing the G95R substitution expressed in bacteria did not significantly change *in vitro* autophosphorylation at multiple regulatory sites, including the activating residue T170, or its specific activity towards a synthetic fluorescent PLK4 peptide substrate (Supplementary Fig. 1C). However, as expected, we observed a marked reduction in inhibition of the G95R PLK4 protein (Supplementary Fig. 1D) by centrinone, and centrinone (and VX-680)-dependent stabilisation of G95R PLK4 protein was decreased by over 75% compared to PLK4 protein, as judged by DSF analysis. These data are consistent with a very significant decrease in PLK4 G95R binding to centrinone, but not Mg-ATP (Supplementary Fig. 1E), which is predicted to manifest as cellular centrinone drug-resistance in the presence of physiological levels of competing ATP.

**Figure 1.**
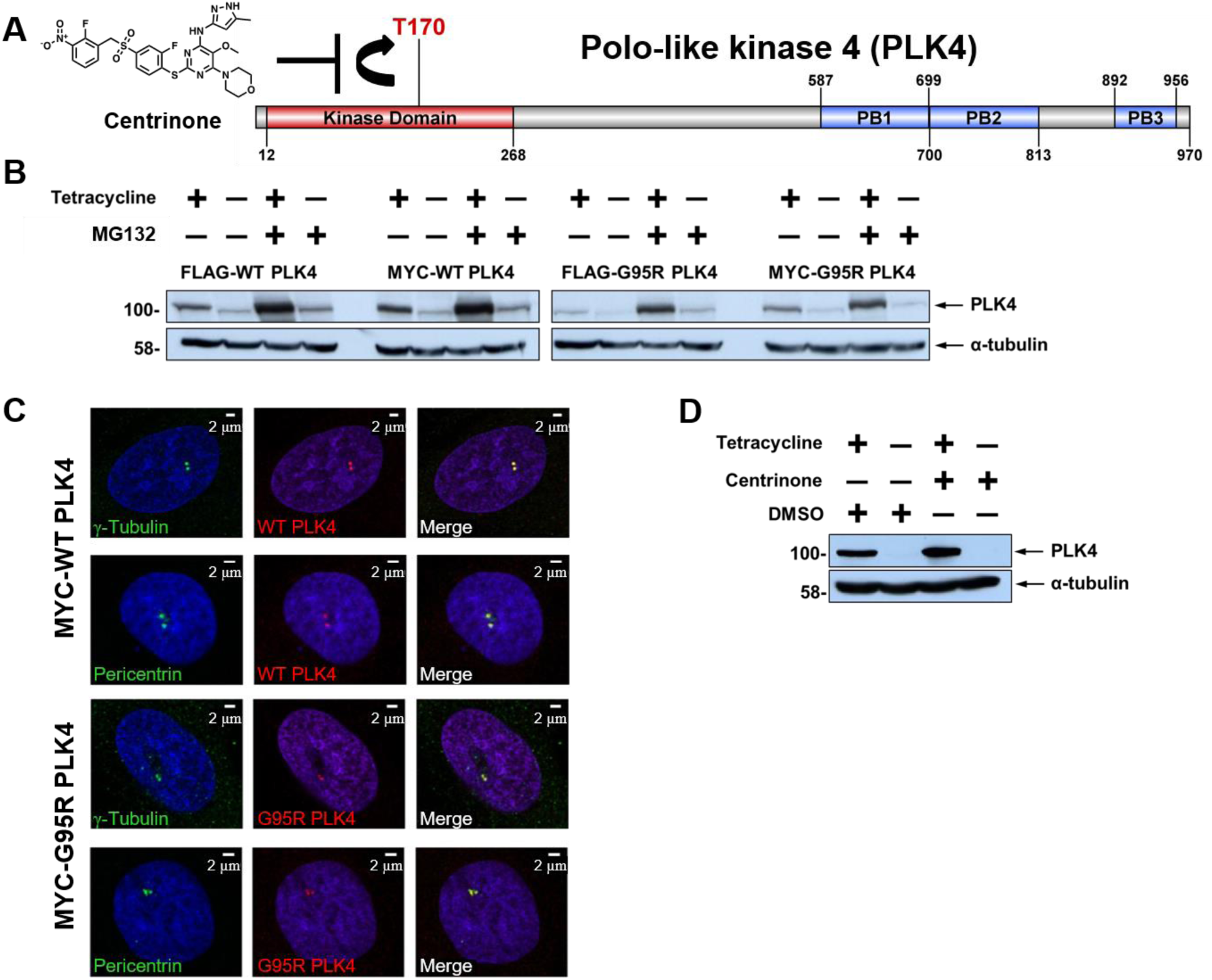
Generation and characterisation of stable inducible cell lines expressing WT or G95R PLK4. **(A)** Structure of the PLK4 inhibitor centrinone and schematic of human PLK4 domain structure. Residues that demarcate the kinase domain, T170 site of activating phosphorylation, and the three polo-box domains are indicated. **(B)** U2OS cells stably transfected with either FLAG- or MYC-tagged WT or G95R PLK4 were incubated in the presence of absence of tetracycline (1 μg/mL) for 18 hours to induce PLK4 expression, followed by MG132 (10 μM) for 4 hours. Lysates were probed with antibodies that recognise either PLK4 or α-tubulin as a loading control. **(C)** U2OS cells induced to express MYC-PLK4 (WT or G95R) with tetracycline were probed with anti-PLK4 (red) and either anti-γ-tubulin or anti-pericentrin (green) antibodies and analysed by immunofluorescence microscopy, confirming localisation of both WT and G95R PLK4 at the centrosome. **(D)** U2OS cells stably transfected with FLAG-WT PLK4 were incubated in the presence or absence of tetracycline (1 μg/mL) for 18 hours. Cells were then treated with 300 nM centrinone or DMSO (0.1% v/v) for 4 hours. Lysates were analysed by western blot and probed with antibodies against either PLK4 or α-tubulin as a loading control.

We next validated inducible expression in WT and G95R PLK4 stable cell lines following incubation with tetracycline, in the absence and presence of the proteasome inhibitor, MG132, which prevents proteasome-mediated degradation of ‘suicide’ kinases such as PLK4 (Fig. 1B; Supplementary Fig. 2). As determined by immunoblotting, both WT and G95R PLK4 expression was induced by tetracycline in all cell lines, with protein levels increasing, as expected, in the presence of MG132. Comparable levels of recombinant PLK4 expression were detected when fused to either an *N*-terminal MYC or FLAG tag (Fig. 1B). By immunofluorescence in fixed cells, we also confirmed that the over-expressed MYC-PLK4 (both WT and G95R) correctly localised to centriolar structures (co-localising with pericentrin and γ-tubulin) during interphase (Fig. 1C). Having generated a new cellular system for controlled over-expression of either WT PLK4 or G95R PLK4, the cellular effects of centrinone inhibition was assessed after Tet-exposure in FLAG-PLK4 lines, which are the focus of the proteomic studies reported below. PLK4 controls its own proteasome-mediated degradation through autophosphorylation of Ser293 and Thr297 in the degron domain [6]; inhibition of PLK4 activity with centrinone should therefore result in PLK4 protein accumulation. As expected, centrinone treatment of stably transfected FLAG-PLK4 increased tetracycline-dependent PLK4 expression levels (Fig. 1D). Owing to the high promiscuity of most ATP-dependent protein kinase inhibitors [73, 74], we next established optimal conditions (inhibitor concentration and length of compound-exposure) for inhibition of PLK4 activity, based on assessment of PLK4 stability, in an attempt to minimise off-target effects. To this end, we exposed WT PLK4 U2OS cells with centrinone and examined FLAG-PLK4 expression at various time points over a 24 h period (Supplementary Fig. 2A). As expected, treatment of the WT PLK4 line with centrinone resulted in time-dependent accumulation of FLAG-PLK4 protein, which was maximal at 300 nM centrinone, similar to MG132 treatment. This observation is consistent with inhibition of PLK4 auto-phosphorylation sites that promote SCF-dependent proteasomal degradation [6]. In contrast, centrinone treatment of FLAG-G95R PLK4 U2OS cells did not result in a marked increase in exogenous PLK4 protein, even at the highest concentration (1 μM) of centrinone tested (Supplementary Fig. 2B, right panel). These cell models therefore provided a centrinone-regulated PLK4 expressing system, which can be used to potentially identify and evaluate PLK4-dependent phosphorylation events and associated signaling pathways.

**Figure 2.**
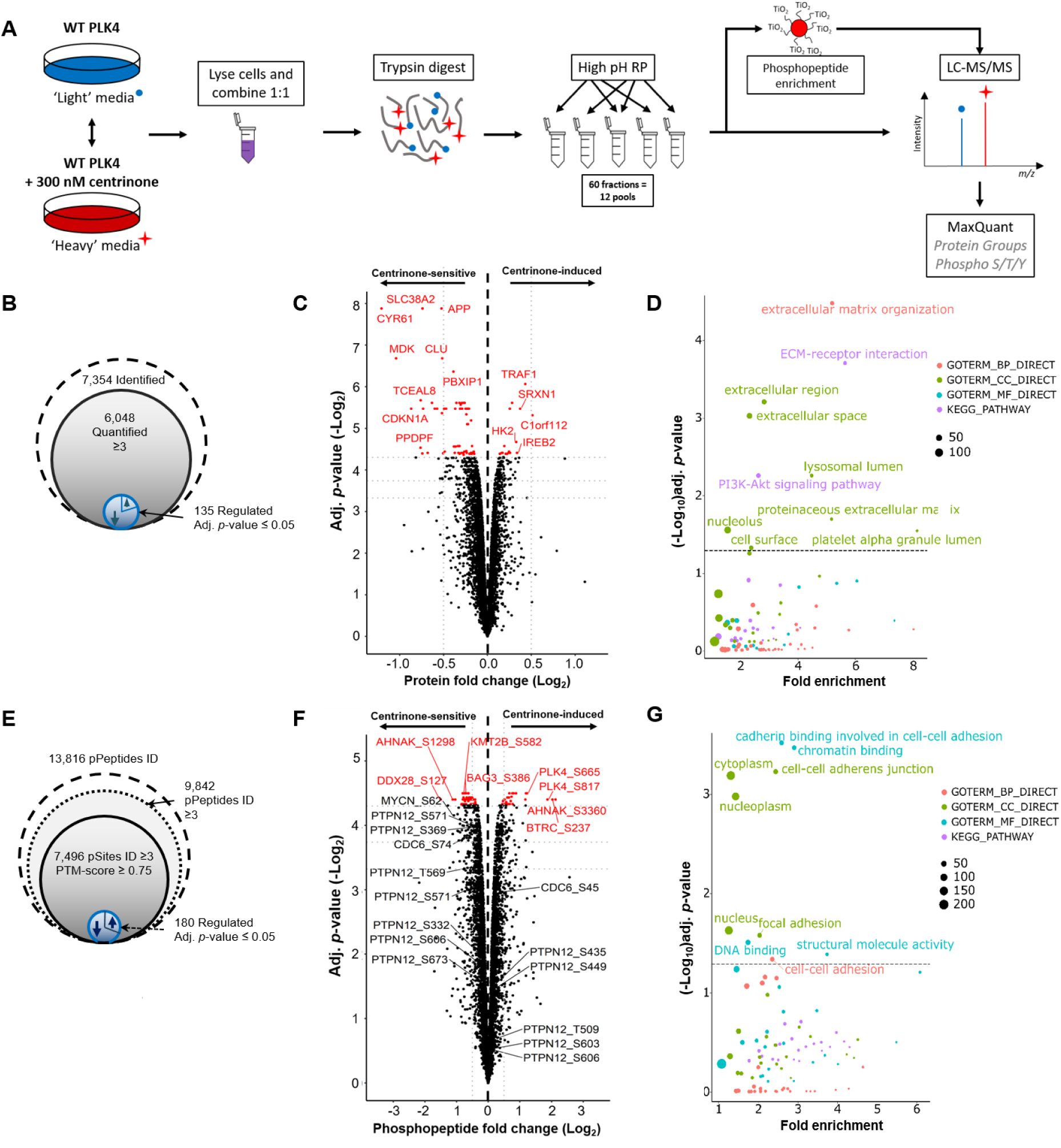
Identification of centrinone-regulated changes in the proteome and phosphoproteome of human cells. (**A**) Workflow for SILAC-based quantitative analysis of centrinone-mediated regulation of the proteome and the phosphoproteome of FLAG-WT PLK4 U2OS cells. Following induction of FLAG-WT PLK4 with tetracycline (1 μg/mL) for 18 hours, cells were treated with either 300 nM centrinone (‘heavy’ labelled) or DMSO (0.1% v/v) control (unlabelled) for 4 hours, prior to protein extraction and tryptic proteolysis. Peptides were separated by high pH reversed-phase chromatography into 12 pools, concatenated from 60 fractions. 90% of each pool was subjected to TiO_2_-based phosphopeptide enrichment. Enriched and non-enriched samples were then subjected to LC-MS/MS using an Orbitrap Fusion mass spectrometer. Data were analysed using MaxQuant software. Total numbers of identified, quantified and differentially regulated (**B**) proteins or (**E**) phosphosites are indicated. Volcano plots showing (**C**) protein or (**F**) phosphopeptide fold ratios following Bayesian statistical analysis to evaluate significant differences. Log_2_-fold change (Heavy/Light) are presented as a function of the –Log_2_ Benjamini-Hochberg adjusted p-value; those with an adjusted p-value ≤0.05 are highlighted in red. Select data points of note are annotated with their protein accession number (and differentially regulated phosphosite in the case of phosphopeptide analysis). (**D, G**) GO term enrichment analysis of significantly regulated proteins (**D**) or phosphosites (**G**) using DAVID. Proteins/phosphopeptides with a Benjamini-Hochberg adjusted p-value ≤0.05 are labelled. BP = biological process (red); CC = cellular compartment (green); MF = molecular function (cyan); KEGG pathway (purple). The size of the node is representative of the number of proteins contributing to a select category.

### SILAC-based centrinone (phospho)proteomics screen

To evaluate centrinone-dependent signaling, and help discover potential new PLK4 substrates, we undertook a global quantitative phosphoproteomics screen using the inducible FLAG-WT PLK4 and FLAG-G95R PLK4 U2OS cell lines. Similar global (phospho)proteomics analyses in the presence of kinase inhibitors have been performed previously to evaluate the cellular activities of PLK1 and Aurora A, revealing phosphorylation sites linked to activity during mitotic progression in HeLa cells [75, 76]. Using a SILAC-based quantification strategy, we evaluated protein and phosphopeptide level regulation following tetracycline induction of FLAG-PLK4 in the absence or presence of centrinone, but in the absence of experimental cell-cycle arrest (Fig. 2A). The efficiency of protein metabolic labelling using ‘heavy’ (R10K8) SILAC media was determined after each cell doubling, and was found to exceed 96%. We also confirmed a lack of significant metabolic conversion [77] of isotopically labelled Arg to Pro (Supplementary Fig. 3). Cells expressing either WT PLK4 or G95R PLK4 were exposed to compounds and harvested after ~7 cell doublings, and lysates for a given cell line were combined for tryptic proteolysis. To improve the depth of coverage of the (phospho)proteome, tryptic peptides were subjected to high-pH reversed-phase chromatography, collecting 60 fractions which were concatenated into 12 pools. A proportion (90%) of these pooled fractions subsequently underwent TiO_2_-based phosphopeptide enrichment and LC-MS/MS analysis, retaining the remaining portion of each sample for comparative total protein quantification (Fig. 2A). A high degree of overlap was observed across the four biological replicates, with 96% of the 7,354 expressed gene products identified in the FLAG-WT PLK4 cell line (at a false discovery rate (FDR) of <1%) being observed in three or more replicates, meaning that a total of 6,046 (82%) proteins could be quantified (Fig. 2B). Not unexpectedly, due to the stochastic nature of data-dependent acquisition (DDA), reproducibility at the phosphopeptide level was lower, with ~71% of the 13,816 phosphopeptides identified in the WT-PLK4 cells being observed in at least three of the biological replicates. Of these, 7,496 sites of phosphorylation, localised at a PTM-score ≥0.75, were quantified across at least three bioreplicates (Fig. 2E).

**Figure 3.**
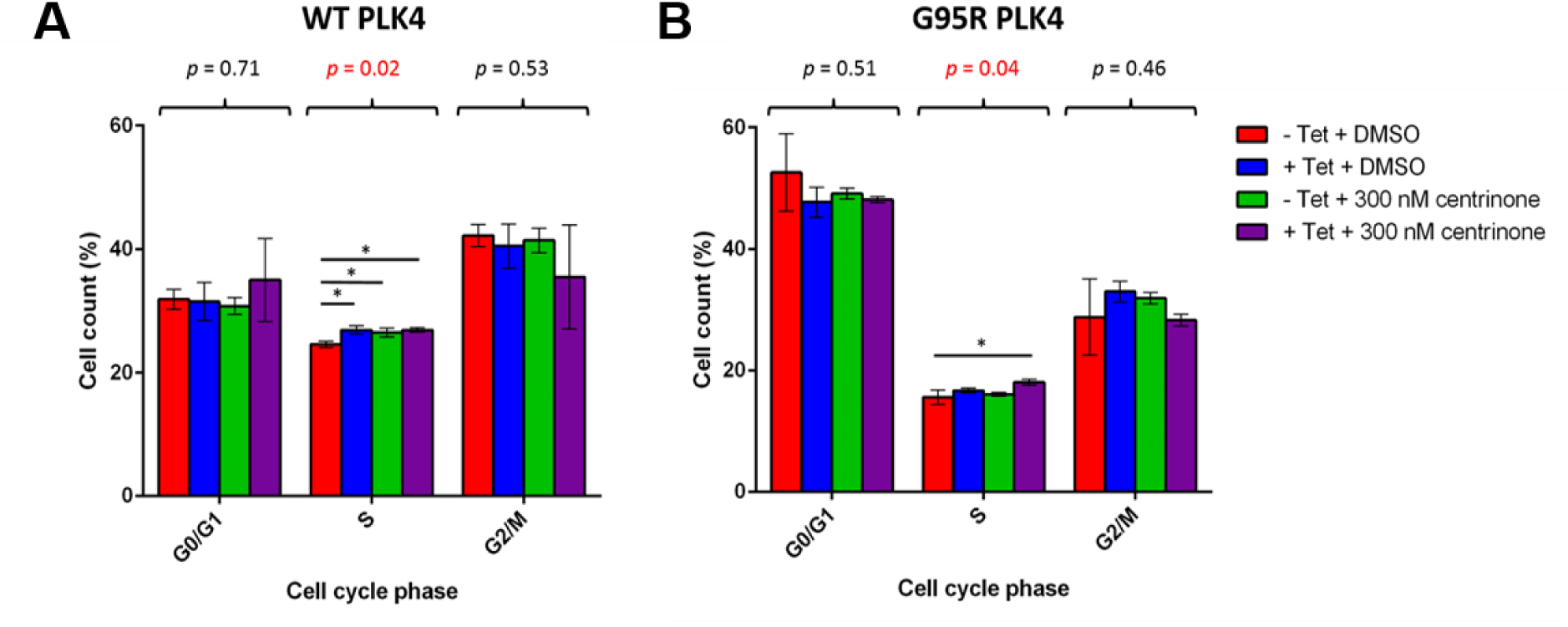
Centrinone induces a subtle S-phase delay in both WT and G95R PLK4 U2OS cells. Expression of FLAG-PLK4 was induced upon incubation with 1 μg/mL tetracycline for 18 hours. Cells were treated with 300 nM centrinone or 0.1 % (v/v) DMSO for 4 hours and cell cycle distribution analysed by FACs. The bar chart reflects the percentage of cells (mean ±SD, n=3) in each cell cycle phase for WT (**A**) and G95R (**B**) PLK4. ANOVA generated p values are shown above each phase. * indicates significant differences between individual samples determined using Tukey post-hoc testing (adj. p value ≤0.05).

At the proteome level, 135 proteins were significantly up or down-regulated upon centrinone treatment of FLAG-WT PLK4 cells, using an adjusted *p-*value ≤0.05, with the level of 100 proteins decreasing, compared to 35 increasing, in the presence of centrinone (Fig. 2B/C and Supplementary Table 1). Importantly, PLK4 itself was upregulated 1.8-fold (*p-*value = 1.0E-03, adj. *p-*value = 0.05) in the presence of centrinone, consistent with accumulation of PLK4 under these conditions, consistent with immunoblotting (Fig. 1D). PLK4 was the only protein that was significantly upregulated (*p-*value ≤0.05) with a greater than 1.5-fold change, although 15 proteins were downregulated (>1.5-fold) under the same conditions, including cyclin-dependent kinase inhibitor 1 (1.8-fold; *p*-value=1.3E-04, adj. *p*-value=2.3E-02) and its binding partner cyclin D2 (1.7-fold; *p*-value=5.8E-04, adj. *p*-value=4.8E-02). Consistent downregulation of the cyclin D2 regulator, CDK4 (1.14-fold, *p*-value=1.1E-03, adj. *p*-value=5.3E-02) was also observed, and the related protein CDK6 was also upregulated under the same conditions (1.16-fold, *p*-value=6.4E-04, adj. *p*-value=4.8E-02). These findings are representative of the several new centrinone targets that associated with G1/S phases of the cell cycle, downstream of PLK4 inhibition. Functional annotation and enrichment analysis of the centrinone-mediated changes in protein abundance using DAVID [67, 68] (Fig. 2D; Supplementary Fig. 4) revealed differential regulation of proteins in a number of distinct cellular compartments including the extracellular region, lysosomal lumen and nucleolus. Proteins involved in extracellular matrix organisation and ECM-receptor interactions were notably down-regulated following centrinone treatment, while nucleolar proteins and those involved in RNA processing/binding were significantly up-regulated (Supplementary Fig. 4). Surprisingly, many more proteins were dysregulated after centrinone treatment in the G95R PLK4 cell line when compared with the WT PLK4-expressing cells (Supplementary Fig. 5). Of the 5,949 proteins quantified in the FLAG-G95R PLK4-expressing cells after centrinone exposure, 1,882 were significantly changed (at an adj. *p-*value ≤0.05). Of these, 585 were upregulated, while 1,297 proteins were down-regulated at the expression level. Functional annotation and enrichment analysis of the centrinone-mediated changes in protein abundance using DAVID revealed differential regulation of proteins localised in the mitochondrion, at focal adhesions and within the cell-cell adherens junctions (Supplementary Fig. 5). The GO term “metabolic pathway proteins”, specifically those involved in amino acid biosynthesis and ribsosome biogenesis, was also enriched, suggesting initiation of a cellular stress response. Proteins involved in cytokinesis and metaphase plate progression were also dysregulated.

**Figure 4.**
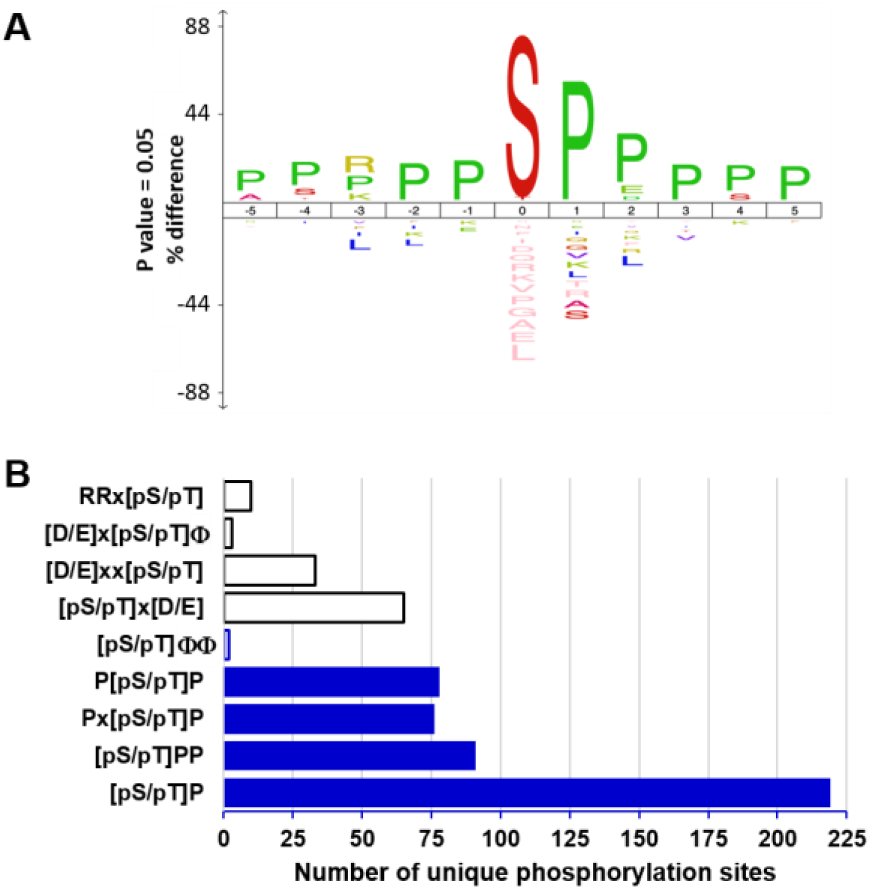
PLK4 phosphorylates serine and threonine residues in a proline consensus. (**A**) Centrinone-down-regulated phosphorylation site sequence conservation using IceLogo [66] reveals extensive Pro enrichment around the site of phosphorylation. (**B**) Frequency of novel Pro-directed motifs (blue), the known consensus sequence for Aurora A (RR[pS/pT]), or the acidic residue-rich motifs of PLK1 ([D/E]x[pS/pT]Φ, where Φ represents hydrophobic Y/F/I/L/V residues and x is any amino acid), PLK2 ([D/E]xx[pS/pT], PLK3 ([pS/pT]x[D/E]) or the classic hydrophobic PLK4 consensus ([pS/pT]ΦΦ) across all centrinone-downregulated phosphorylation sites.

**Figure 5.**
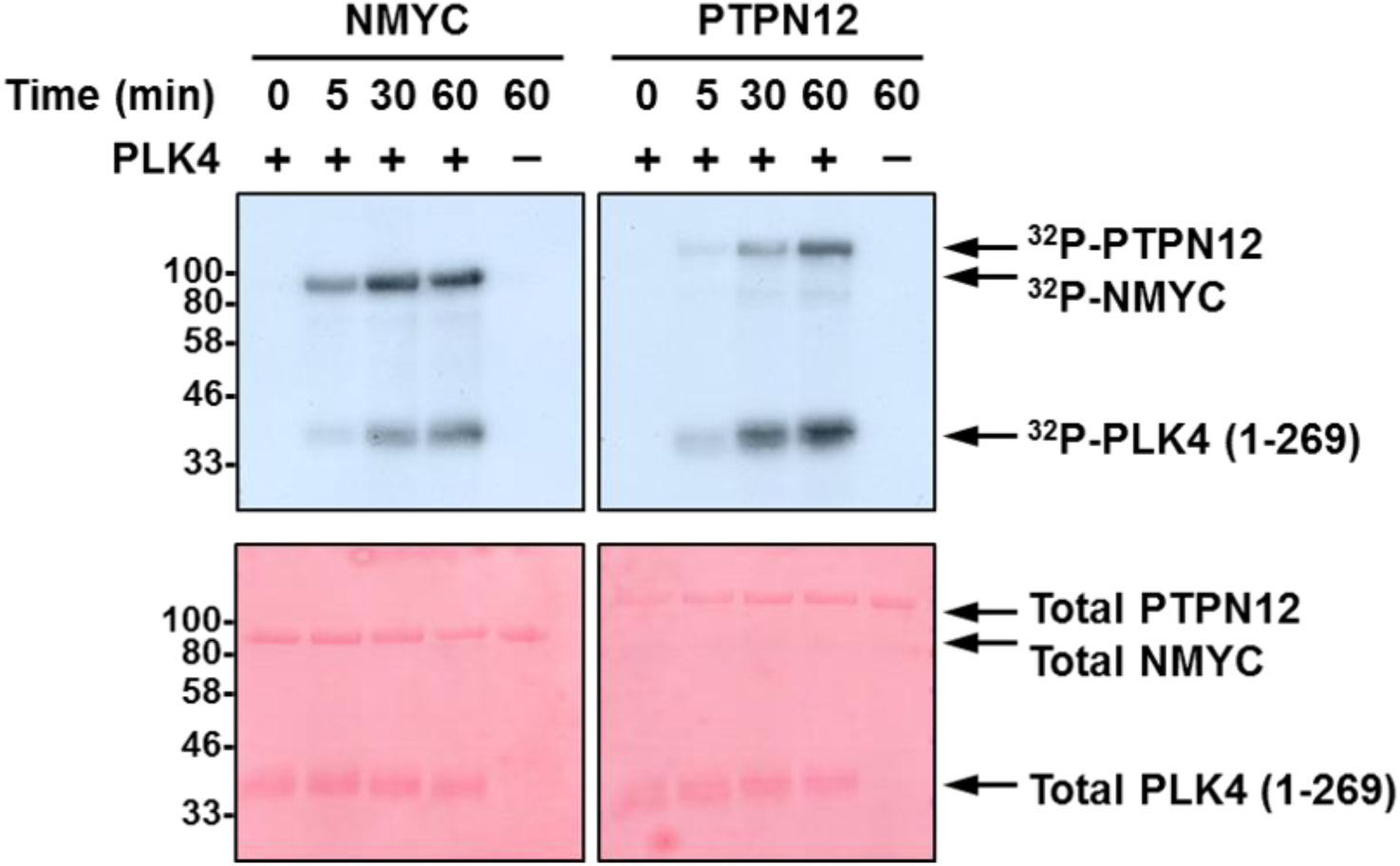
PLK4 phosphorylates NMYC and PTPN12 *in vitro*. Autoradiogram (top) or Ponceau stained membrane (bottom) following in vitro ^32^P-based phosphorylation of either recombinant NMYC or PTPN12 by recombinant PLK4 catalytic domain.

Interestingly, several known mitotic proteins were significantly down-regulated in G95R PLK4-expressing cells, including Aurora A (1.6-fold, *p*-value=6.0E-05, adj. *p*-value=4.9E-03), TPX2 (1.6-fold, *p*-value=7.7E-06, adj. *p*-value=3.5E-03), cyclin B1 (1.6-fold, *p*-value=9.2E-07, adj. *p*-value=2.7E-03), cyclin B2 (1.4-fold, *p*-value=4.2E-03, adj. *p*-value=2.4E-03) and FOXM1 (1.5-fold, *p*-value=6.1E-06, adj. *p*-value=3.0E-03), suggesting centrinone effects on a G2/M regulatory protein network. Depletion of sororin (1.6-fold, *p*-value=3.1E-06, adj. *p*-value=3.0E-03) and CDC20 (1.6-fold, *p*-value=8.9E-06, adj. *p*-value=3.8E-03), key regulators of sister chromatid cohesion, and borealin (CDCA8; 1.5-fold, *p*-value=1.1E-05, adj. *p*-value=3.8E-03), a component of the mitotic CPC complex, also implicates a negative effect of centrinone on chromosome cohesion in clonal G95R PLK4 cells. Functional network analysis using STRING [69] revealed a broader connection between these down-regulated G2/M-phase proteins (Supplementary Fig. 5D) suggesting that centrinone exposure targeted pathways controlling a broad spectrum of cell cycle regulators. Interestingly, many of those were not identified in WT PLK4-expressing cells, consistent with the presence of newly-engaged non-PLK4 “off-targets” (including several linked to Aurora A and G2/M phase) in cells expressing the centrinone-resistant PLK4 mutant.

Flow cytometry-based analysis of the cell cycle distribution of U2OS cells following induction of either WT PLK4 or G95R PLK did not reveal significant changes in G1 or G2/M cell cycle distribution (ruling-out induced kinase activities that are found in these phases), although both tetracycline (which leads to increased WT and G95R PLK4 expression) and centrinone (chemical PLK4 inhibition) induced subtle S-phase enrichment (Fig. 3). Overall, these data are consistent with the protein-level changes observed in both cell lines, with a small increase in S-phase cells suggesting either a slight S-phase delay, or the early onset of mitotic arrest. Further work is needed to tease apart the ‘on’ (PLK4) and ‘off’ (non-PLK4) targets of centrinone in cells that become engaged under these conditions. However, the comparatively few proteins whose levels are regulated upon centrinone treatment in the WT PLK4 cell line (~1% of total) compared with the G95R PLK4 cell line (~30% of total), combined with our chemical genetic confirmation of centrinone as an on-target PLK4 inhibitor, leads to an interesting observation; in G95R PLK4 cells in the absence of its ‘preferential’ exogenous binding partner WT-PLK4, centrinone also induces destabilisation and dephosphorylation of multiple off-targets. We hypothesise (but are not yet able to prove) that this could reflect the ability of the compound to inhibit a low affinity ‘off-target’ such as Aurora A, when binding to the higher affinity target protein PLK4 is compromised.

### The centrinone-modulated phosphoproteome

Of the ~7,500 ‘class I’ phosphosites (PTM-score ≥0.75) identified in clonal WT PLK4 cells (Fig. 2E, F), 183 (~2.4%) were on peptides that were differentially regulated by centrinone using an adjusted *p*-value ≤0.05 (Supplementary Table 2). Of these, 130 phosphorylation sites were localised with ≥99% site localisation confidence, *i.e.* PTM-score ≥0.994 [60]. The vast majority of the statistically differentially regulated phosphopeptides (containing 135 ‘class I’ phosphosites, 98 of which were localised at a 1% FLR) were down-regulated, consistent with inhibition of at least one protein kinase, or activation of at least one protein phosphatase. Using a slightly less stringent adjusted *p*-value ≤0.075 (where all *p*-values were ≤0.005), 468 (~6%) phosphopeptides containing 480 phosphosites (348 at 1% FLR) were differentially regulated by centrinone-exposure, 318 of which were reduced, and might be considered to be potential PLK4 substrates.

To understand how many of these regulated phosphosites were considered to be previously ‘known’, we mapped them against data from large scale data repositories: PeptideAtlas (PA) phosphopeptide builds, and PhosphoSitePlus (PSP), categorizing data from the different sources according to the strength of evidence (see Methods). From this analysis, 99% of our identified ‘class I’ sites were considered high or medium confidence sites within PSP, and 94% considered high or medium confidence sites in PA. These comparisons both support our findings that these sites have been correctly localized in the vast majority of cases, and confirm previous studies. Randomly incorrectly localized sites would generally occur with low confidence or fail to have supporting evidence from PSP and/or PA.

We also calculated the evolutionary conservation of the phosphosites across 100 model proteomes (50 mammals, 12 birds, 5 fish, 4 reptiles, 2 amphibians, 11 insects, 4 fungi, 7 plants, 5 protists; Supplementary Table 2), demonstrating that most of the sites are conserved in at least half of the proteomes, allowing for conservative S/T substitutions (median 51%, upper quartile 68%; lower 44%). In the vast majority of cases, there are examples of conservation across mammals, birds, reptiles and fish, suggesting that these phosphorylation sites may contribute broadly to metazoan signaling pathways. Several sites, including Ser192 on Cdc42 effector protein 1 (downregulated 1.6 fold in response to centrinone; *p-*value = 1.8E-03) and Ser48 in histone H4 (downregulated 1.3-fold, *p-*value = 1.60E-03) are ~100% conserved across all eukaryotes tested (Supplementary Table 2).

Three differentially regulated phosphorylation sites were identified on PLK4 itself: Ser421, Ser665 and Ser821 (which are conserved in 55%, 44% and 43% respectively of 94 aligned species) all increased in response to centrinone treatment (Supplementary Fig. 6, Supplementary Table 2). While the fold change for Ser421 matched the change in PLK4 protein level expression (both were up-regulated 1.8-fold), indicating no overall change in phosphorylation stoichiometry at this site, levels of pSer665 and pSer821 (2.3- and 3.7-fold respectively) were notably higher in the presence of centrinone, suggesting phosphorylation of these two sites by a regulatory kinase or phosphatase. Although both Ser665 and Ser821 have previously been demonstrated to be phosphorylated in high-throughput MS-based studies [78], their functional roles remain to be defined. We did not find any evidence for changes in canonical multi-phosphodegron sites of phosphorylation in PLK4, likely due to the rapid turnover of PLK4 that occurs after phosphorylation at these sites.

**Figure 6.**
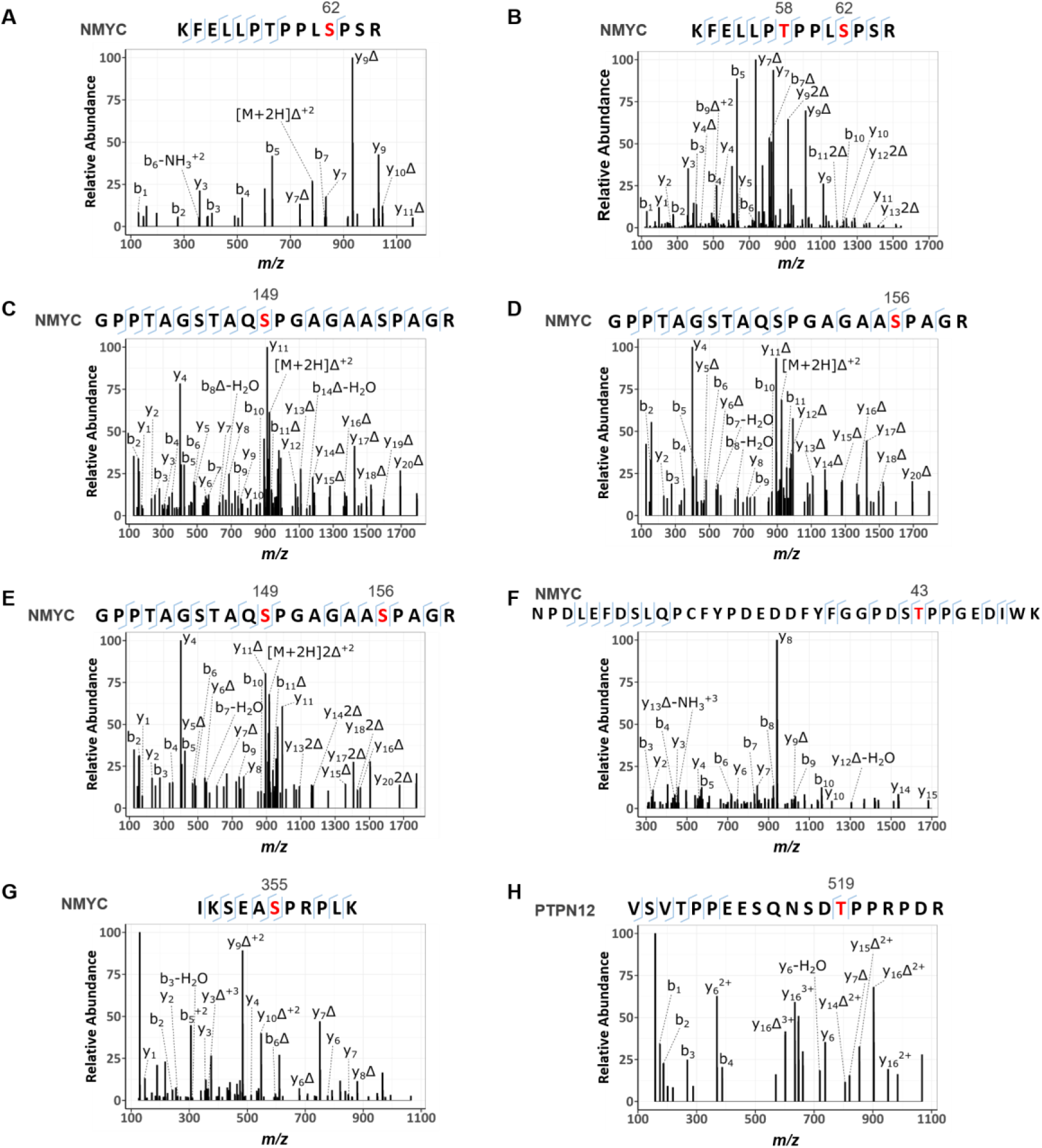
HCD product ion mass spectra of PLK4-dependent [pS/pT]P phosphorylated tryptic peptides from NMYC and PTPN12. Recombinant NMYC **(A-G)** and PTPN12 **(H)** proteins were incubated in 50 mM Tris-HCl, pH 7.4, 100 mM NaCl, 1 mM DTT, 1 mM ATP and 5 mM MgCl_2_ at 30°C in the absence and presence of PLK4. Reactions were then digested with trypsin and subjected to TiO_2_–based phosphopeptide enrichment and LC-MS/MS analysis, as described in Methods. **(A)** doubly charged ion at m/z 831.4367 indicates phosphorylation of NMYC at Ser62; **(B)** triply charged ion at m/z 581.2839 indicates phosphorylation of NMYC at Thr58 and Ser62; **(C, D)** doubly charged ion at m/z 973.4420 indicated phosphorylation of NMYC at either Ser149 **(C)** or Ser156 **(D); (E)** triply charged ion at m/z 675.9525 indicated phosphorylation of NMYC at Ser149 and Ser156**; (F)** triply charged ion at m/z 1430.5940, indicated phosphorylation of NMYC at Thr43; **(G)** triply charged ion at m/z 435.9026 indicates phosphorylation of NMYC at Ser355; **(H)** triply charged ion at m/z 763.3448 indicates phosphorylation of PTPN12 at Thr519. Red denotes the site of phosphorylation. Δ equates to loss of H_3_PO_4_.

Gene ontology (GO) analysis using DAVID also revealed that centrinone-regulated phosphorylation sites (adj. *p*-value ≤0.075) were significantly enriched in proteins involved in chromatin (up-regulated) and cadherin binding (down-regulated) (Fig. 2G; Supplementary Fig. 4). Interestingly, phosphoproteins in the nucleoplasm were also up-regulated, while levels of cytoplasmic proteins were reduced in a centrinone-dependent manner. In total, 22 centrosome phosphoproteins were significantly regulated, with the levels of 8 phosphoproteins increasing and 14 being reduced, including NUMA1 (pSer1769, pSer1991), LZTS2 (pSer311), RANBP1 (pSer60), XRCC4 (pSer256) and WDR62 (pSer33), a target of the physiological PLK4 targets CEP152 and CEP192 [11, 12, 37, 79–81]. Phosphorylation sites on five members of the MAPK signaling pathway were also statistically differentially regulated: pThr693 on EGFR (*p*-value=5.0E-04, adj. *p*-value=5.0E-02) and pTyr204 on ERK1 (*p*-value=3.4E-03, adj. *p*-value=6.8E-02) were both up-regulated 1.5-fold, while pSer1275 on SOS1 (1.6-fold down-regulated, *p*-value=1.1E-03, adj. *p*-value=5.2E-02), pSer23 on MEK2 (MAP2K2; 1.5-fold, *p*-value=1.2E-04, adj. *p*-value=4.6E-02), and pSer62 on NMYC (2.1-fold, *p*-value=1.4E-03, adj. *p*-value=5.6E-02) were all down-regulated at least 1.5-fold. EGFR was the only one of these proteins to be differentially regulated at the protein level (1.2-fold up-regulation), although MAP3K2 (MEKK2), an upstream MAPK pathway regulator, was also down-regulated 1.2-fold at the protein level at an adj. *p*-value=5.5E-02. Consequently, PLK4-dependent signaling is implicated in regulation of the MAPK pathway, which in-turn can control entry into the cell cycle, amongst other functions. Consistent with this, and with the protein level changes discussed above, we observed centrinone-mediated regulation of a number of phosphopeptides from proteins implicated in cell cycle processes, including peptides containing: either i) pSer277/pSer283 or ii) pSer283/pSer285 from CDK11B (both down 1.3-fold); iii) pSer332/pSer334 (1.2-fold up), (iv) pSer644 (1.3-fold down), (v) pSer681/pSer685 (1.5-fold down) on CDK12; (vi) pSer130 on CDK18 (1.2-fold down); (vii) pSer37 (1.4-fold down) and (viii) pThr821/826 (1.5-fold up) on RB1; (ix) pSer271 on cyclin D2 (2-fold down); (x) pSer387 on cyclin E1 (1.3-fold down). Interestingly, pSer74 on CDC6 was also reduced 1.3-fold in the presence of centrinone. CDC6 and PLK4 work antagonistically to regulate centriole duplication; this is driven by PLK4-mediated disruption of the CDC6-Sas-6 complex, and thus the ability of Sas-6 to interact with the centriole duplication core protein STIL. PLK4 binds directly to the *N*-terminal region of CDC6 in S-phase, disrupting the CDC6 Sas-6 interaction following CDC6 phosphorylation. Based on prediction and mutational analysis, PLK4 was previously shown to phosphorylate CDC6 *in vitro* on Ser30 and Thr527 [82]. Based on our data, we suggest that Ser74, which is conserved in ~70% of aligned eukaryotic orthologues, is a potential additional phosphosite on CDC6 that lies in a SerPro consensus downstream of centrinone in cells [82]. In agreement with the hypothesis that this is a direct PLK4 target, this phosphopeptide was not observed as differentially-regulated in the G95R PLK4 cell line when treated with centrinone.

From the limited selection of physiological PLK4 substrates identified to date [38], we did not identify any known phosphopeptides that were statistically downregulated in response to centrinone, likely due to our experimental procedure, in which cells were not chemically synchronised in S-phase despite centrinone-induced PLK4 stabilisation. Surprisingly, we observed a greater than 4-fold increase in levels of the phosphopeptide containing pSer237 (with respect to protein-level changes) of BTRC (F-box/WD repeat-containing protein 1A), which has been previously reported to be associated with the PLK4 activator STIL [83], and 1.3-fold up-regulation of pSer3467 on the E3 ubiquitin-protein ligase MYCBP2.

Chemical genetic PLK4 strategies have previously reported decreased phosphorylation of multiple proteins, including RUNX1, PTPN12, IL6ST, TRIM3 and SCRIB after conditional PLK4 knockdown or transgenic expression and inhibition of ‘analogue-sensitive’ PLK4 [13, 35, 38, 84]. Many of these proteins have been assumed to be indirect targets of PLK4, due to the fact that they are not known to localise to the centrioles (or play known roles in centrosome or cilia biology). In our datasets, we also identified a number of confidently-localised phosphosites for most of these proteins, notably Ser369 and Ser571 on PTPN12, a purported ERK site of phosphorylation [85], and Ser1309 and Ser1348 on SCRIB which were statistically down-regulated (*q*-value ≤0.075) after centrinone exposure, suggesting simplistically that they are likely to lie downstream of PLK4. Considering a reduced confidence threshold of statistical regulation (*p*-value <0.05), Thr569 phosphorylation on PTPN12, as well as Ser708 and Ser1306 phosphorylation on SCRIB, were also found to be significantly down-regulated in our datasets. The use of drug-resistant kinase alleles can efficiently confirm on-target effects of small molecules [48, 55, 56]. However in the case of drug-resistant G95R PLK4, Tet-induced cells behaved differently to WT-PLK4 in terms of effects on known PLK4 phosphorylation site motifs, meaning that we could not use experimental cellular evidence to validate on-target PLK4 phosphorylation with this system. Consequently, we resorted to biochemical methods.

### Can PLK4 directly phosphorylate a Pro-rich amino acid consensus in proteins?

Motif analysis of the 318 phosphorylation sites on peptides that were down-regulated in response to centrinone (adj. *p*-value ≤0.075) revealed very strong enrichment for phosphorylation sites in a Pro-rich consensus, with a Pro residue prevalent at the +1 position (Fig. 4A; Supplementary Table 2). Indeed, 219 (69%) of the unique (type I) phosphosites identified in cells were immediately followed by a Pro residue (Fig. 4B), with 91 phosphorylation sites additionally containing Pro at +2 ([pS/pT]PP consensus), while 78 and 76 phosphosites contain a Pro at −1 or −2, respectively (P[pS/pT]P or Px[pS/pT]P). Although a canonical PLK4 peptide phosphorylation consensus motif has been described in which one or two hydrophobic (Φ) residues reside immediately *C*-terminal to the site of phosphorylation [47], only 2 (~1%) of the identified centrinone-down-regulated phosphorylated sites contained Y/F/I/L/V in both of these ‘canonical’ positions (Fig. 4B). Even considering a single hydrophobic residue at either positions +1 or +2, the number of phosphosites within this motif was significantly lower at 52, than the number observed within a Pro-rich consensus (216, Fig. 4B). Interestingly, 2 of the 16 autophosphorylation sites that we identified on recombinant PLK4 (Ser174 and Ser179) were also followed by a Pro residue (Supplementary Fig. 1B), indicating that a Pro at this position is permissible for phosphorylation by PLK4. In considering potential off-target effects of centrinone on either Aurora A or other PLK family members, we also interrogated the down-regulated phosphorylation sites for the RRx[S/T] Aurora A consensus and the classic [D/E]x[S/T]Φ consensus of PLK1 [42]. However, only 10 centrinone inhibited sites were observed that contained a basic Arg residue at both −1 and −2, and only three sites fitted the PLK1 substrate consensus (Fig. 4B). In agreement with these observations, we also determined that a significant proportion (47%) of the 141 phosphopeptides downregulated in a recent study evaluating PLK4 substrates using analogue-sensitive alleles in human RPE-1 cells [38] also contained a Pro at +1 relative to the site of modification, compared with only 15% that possess a hydrophobic residue at +1 and/or +2 from the site of phosphorylation.

To experimentally validate centrinone-regulated sites identified as potential PLK4 substrates in cells, and to examine our hypothesis of a potential Pro-driven consensus motif for this kinase, several recombinant proteins were evaluated for their ability to be phosphorylated by active PLK4 *in vitro.* Both NMYC and PTPN12 contained phosphorylation sites with a Pro at +1 that were statistically down-regulated following cellular exposure to centrinone: pSer62 on NMYC (2.1-fold, *p*-value=1.4E-03, adj. *p*-value=5.6E-02, conserved in 71% of aligned species), a site previously reported to be phosphorylated by cyclin B/Cdk1 in prophase [86], and two sites on PTPN12: pSer369 (1.3-fold, *p*-value=2.4E-03, adj. *p*-value=6.4E-02, conserved in 89% of species) and pSer571 (1.3-fold, *p*-value=2.4E-03, adj. *p*-value=6.4E-02, conserved in 60%), a site previously shown to be phosphorylated by ERK1/2 *in vitro* [85]. Interestingly, both PTPN12 phosphosites also contained a Pro at −1 as well as a Pro at +1. NMYC and PTPN12 were thus selected as potential PLK4 substrates for biochemical analysis. PLK4 kinase assays employing either recombinant NMYC or PTPN12 as substrates readily confirmed efficient (similar to PLK4 autophosphorylation) phosphorylation of both proteins *in vitro* (Fig. 5, 6; Table 2). MS-based phosphorylation site mapping of recombinant NMYC with catalytically-active recombinant PLK4 confirmed Ser26(Pro) as a PLK4-dependent phosphorylation site (Table 2; Fig. 6). A number of additional PLK4-dependent phosphosites were also identified, including <u>Thr43</u>, <u>Thr58</u>, Ser131, <u>Ser149</u>, <u>Ser156</u>, Ser315, <u>Ser355</u> and Ser400, five of which (underlined) also possess a Pro at the +1 position. Interestingly, this region of NMYC contains an extended Aurora A docking site in the N-terminus [87], which also hosts a phosphodegron centred around Pro-directed Thr58, whose phosphorylation via GSK3 and a Pro-directed kinase has been shown to be required for recognition by the E3 ubiquitin ligase SCF^FbxW7^ [88, 89]. In a separate experiment, full-length recombinant PTPN12 was also phosphorylated by PLK4 *in vitro* on four residues (Ser435, Ser499, <u>Thr519</u>, Ser606), including notably Thr519, which lies in a pTPP consensus (Table 2, Fig. 6). Ser369 and Ser571, which changed in cells in the presence of centrinone, were not identified in under these conditions (Table 2).

### Analysis of peptide phosphorylation by recombinant PLK4

To further evaluate the hypothesis that PLK4 might function as a Pro-directed kinase, we undertook a series of peptide-based kinase assays beginning with our standard PLK4 peptide substrate, FLAKSFGSPNRAYKK, which is derived from the activation segment of CDK7/CAK [27]. This peptide contains two potential sites of Ser/Thr phosphorylation, including one Ser-Pro site. The precise site of PLK4-dependent phosphorylation in this peptide substrate has not previously been defined. Based on the quantitative phosphoproteomics data, and the protein assays described above, we hypothesised that PLK4 directly catalyses phosphorylation of Ser8 in this peptide, since this phosphoacceptor residue is immediately followed by Pro.

However, MS-based phosphosite mapping revealed that the incorporated phosphate quantified by microfluidic phosphopeptide mobility assay (Fig. 7A) resides solely on Ser5, as opposed to Ser8, of this peptide (Supplementary Fig. 7). Phosphorylation of Ser5 was then confirmed (indirectly) by using a variant peptide where Ser5 was replaced with Ala (S5A), which was not phosphorylated by PLK4 (Fig. 7A). Interestingly, the single mutation of Pro9 to Ala also greatly reduced phosphorylation at Ser5, through an unknown mechanism that presumably involves peptide recognition to prime Ser8 phosphorylation. To supplement this analysis, a different PLK4 peptide substrate lacking a Pro residue was synthesised for mobility-based kinase assays based on the published canonical [pS/pT]ΦΦ PLK4 phosphorylation consensus motif, (RKKKSFYFKKHHH) termed ‘RK’ [47]. This peptide was also exploited to evaluate the effects of a Pro at +1 by single amino acid replacement (RKKKS<u>P</u>YPKKHHH). The substrate was phosphorylated more efficiently by PLK4 than our standard peptide (Fig. 7B), but replacement of Phe at +1 with a Pro residue completed abolished PLK4-dependent phosphorylation, as previously reported in high-throughput studies [47]. Thus, although Pro at +1 is permissible for direct PLK4 phosphorylation in some protein substrates (Fig. 1, Table 2, Fig. 6), synthetic peptide data demonstrate that Pro at the +1 position is not tolerated when short peptide motifs are presented as potential substrates, despite evidence for phosphorylation in intact proteins containing these sequences.

**Figure 7.**
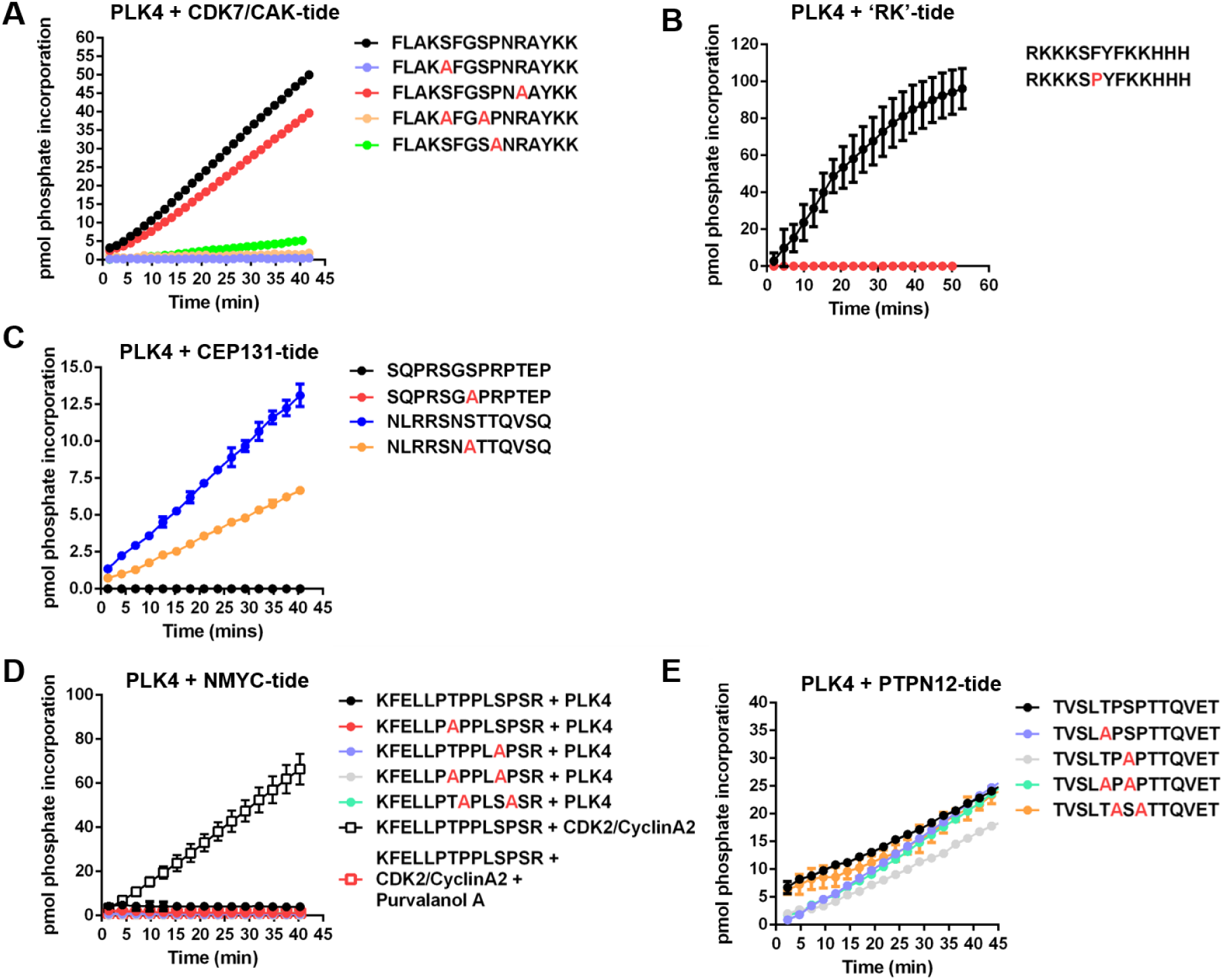
PLK4 substrate specificity differs towards peptide substrates. PLK4 or CDK2/CyclinA2 activity was quantified in a time-dependent manner using the indicated fluorescently-labelled synthetic peptide substrates: (**A**) CDK7/CAK derived substrates; (**B**) ‘RK’ and ‘RK’ F6P substrate; (**C**) CEP131 derived substrates; (**D**) NMYC derived substrates; (**E**) PTPN12 derived substrates. All assays were performed at room temperature (20 °C) using 2 μM final concentration of the appropriate peptide substrate, 1 mM ATP and 500 ng PLK4 or 10 ng CDK2/CyclinA2 as indicated. The specific activity (pmol phosphate incorporation) was calculated from integrated phospho:dephosphopeptide ratios at the indicated time points after assay initiation.

Next, we turned our attention to PLK4 substrates derived from known cellular substrates, including those lying downstream of centrinone. Initially, we evaluated two (overlapping) peptides derived from phosphorylated CEP131, a centriolar PLK4 substrate [38]. We confirmed that peptides containing the sequence surrounding Ser78 (in an N<u>S</u>TT motif), but surrounding Ser89 (lying in a G<u>S</u>PRP motif), were phosphorylated in real-time by PLK4. The substitution of Ser78 for Ala reduced phosphorylation by approximately 50%, suggesting additional peptide modification site(s) in this peptide (Fig. 7C). We also evaluated a peptide substrate designed around the Thr58/Ser62 [pS/pT]P phosphorylation sites that we identified in NMYC, which are centrinone-dependent events in cells (pSer62; Supplementary Table 2) and PLK4 phosphorylation sites in NMYC protein *in vitro* (Fig. 6B; Table 2). Surprisingly, none of the synthetic short NMYC peptides were detectably phosphorylated by PLK4 (Fig. 7D). Moreover, to confirm the ability of this peptide to function as a kinase substrate we showed that it was phosphorylated by the Pro-directed kinase CDK2/cyclinA2 in a Purvalanol-dependent manner (Fig. 7D). Finally, although a peptide designed around the cellular pT569/pS571 phosphosites identified in PTPN12 (which are centrinone-responsive) was a substrate for PLK4, this was largely independent of a Pro residue at the +1 position at either of these residues, based on similar rates of phosphorylation after mutation of either or both of these phosphorylatable residues to Ala (Fig. 7E). This is in broad agreement with our *in vitro* protein data for PTPN12 (Table 2), where we found that although recombinant PTPN12 is a PLK4 substrate (including a single validated pTP consensus), centrinone-sensitive Pro-directed PTPN12 phosphorylation sites identified in cells were not detectably phosphorylated *in vitro*.

## DISCUSSION

In this study, we developed a quantitative phosphoproteomics approach to explore PLK4-mediated signaling in human cells and attempted to identify novel cellular PLK4 targets by exploiting the target-validated PLK4-selective inhibitor centrinone. Of the 480 phosphorylation sites identified on peptides that were differentially regulated in FLAG-WT PLK4 U2OS cells exposed to centrinone (at an adjusted *p*-value ≤0.075, where all *p*-values were ≤0.005), 318 were found on peptides whose levels decreased in the presence of the compound, suggesting either direct (or indirect) regulation through inhibition of PLK4 catalytic activity. The majority of these sites are conserved across eukaryotes, suggesting evolutionary conserved PLK4-mediated signaling mechanisms. Surprisingly, sequence analysis of the residues surrounding these down-regulated phosphorylation sites revealed a highly dominant Pro-rich phosphorylation consensus, notably at positions +1, −1/-2 with respect to the site of phosphorylation (Fig. 4A). Indeed, ~70% of the phosphosites that were reduced in the presence of centrinone possessed a Pro at +1, with 42% of those also containing a Pro residue at +2 (Fig. 4B).

As well as identifying a number of centrinone-inhibited phosphorylation sites on multiple cell cycle-regulating proteins that might be PLK4 substrates (including CDK11, CDK12, CDK18, RB1, cyclin D2, cyclin E1 and CDC6), we report two new centrinone-regulated proteins as direct PLK4 substrates, NMYC and PTPN12, which we validate using *in vitro* kinase assays and MS-based site analysis (Figs. 5 and 6). Of 13 PLK4-dependent phosphorylation sites mapped in NMYC and PTPN12 (9 and 4 respectively), 6 are followed by a Pro residue (Table 2, Fig. 6), supporting the idea that in some protein substrates, PLK4 possesses an ability to function as a Pro-directed Ser/Thr kinase. Previous high-throughput studies agree with these findings, including the mapping of Pro-directed autophosphorylation sites on PLK4 [26, 27], and *in vitro* MS-based site-mapping demonstrating PLK4 phosphorylation of Ser and Thr residues followed by a Pro residue in the physiological substrate STIL [16]. These findings contrast with other assays that employ synthetic substrates, including sequence consensus motifs identical to phosphorylation sites mapped in cellular PLK4 targets [38, 46, 47] (Fig. 6, Table 2 and Supplementary Table 2). We thus hypothesise that under some conditions, PLK4 has the ability to phosphorylate [pS/pT]P motifs in a manner that is dependent on long-range interactions within its protein substrates. For example, these could involve phosphorylated Polo box-binding sequences deposited on PLK4 substrates by PLK4 itself, or by a variety of ‘priming’ kinases. Whatever the mechanisms at play, our data suggest that comparison of substrate recognition sequences from peptide-based screening should be used with some caution when evaluating cellular proteomics data to understand kinase substrate relationships. Indeed, in the case of previous phosphoproteomics datasets obtained with analogue-sensitive PLK4 and PP1 analogues, or through *in vitro* mapping of physiological substrates such as Ana2/STIL, multiple [pS/pT]P sites have been deposited in databases, but appear to have been largely ignored. As a specific example, of 84 STIL phosphorylation sites directly phosphorylated by PLK4 *in vitro*, 14 reside in either an pSP or pTP consensus [16]. Moreover, the phosphorylation of putative PLK4 substrates, including RUNX1, TRIM3, SCRIB and CEP131 on [S/T]P sites [38, 90], have been reported, and many of these PLK4-phosphorylated [pS/pT]P motifs are conserved in eukaryotes.

Although we validate NMYC and PTPN12 as new PLK4 substrates, we cannot rule out that some of the (Pro and indeed non Pro-directed) centrinone-inhibited phosphorylation sites found in the cellular proteome are either i) off-target to centrinone, or ii) indirect effects of PLK4 inhibition of a downstream Pro-directed kinase, perhaps one that is involved in ‘priming’ PLK4 binding via one of the PLK4 PBDs. In an attempt to mitigate against such off-target effects of centrinone, we exploited cell lines expressing G95R PLK4 that is predicted to be unable to bind the inhibitor, and remains catalytically viable (Supplementary Fig. 1). However, based on the results reported here, we predict that interpretation of the quantitative phosphoproteomics data with this cell line are confounded by centrinone-mediated inhibition of both the endogenous PLK4 protein, which is still present at low levels in these studies, and ‘off-targets’ of the compound. In considering cellular targets of centrinone, including potential off-targets of the VX-680 parental compound, such as Aurora A and B, we also interrogated the sequence context of downregulated phosphosites for the classical Aurora A/B (and PLK1/3) substrate motifs. Although phosphorylation sites were identified on down-regulated phosphopeptides that matched the basophilic consensus motifs for these kinases, when combined, these still only accounted for <25% of the ‘non-Pro’ directed motifs identified, suggesting that inhibition of these enzymes is not a major confounding issue for analysis of data obtained with this compound. The total depletion of endogenous PLK4 (*e.g.* by CRISPR/Cas9) strategies is challenging due to its fundamental role in cellular biology, thus it has not been possible for us to generate an inducible system in a ‘clean’ PLK4 null background. However, it will be of interest in future to engineer a G95R PLK4 germline mutation and then model specific ‘on’ and ‘off’ targets of centrinone more carefully under a series of defined experimental conditions.

We observed that many Pro-directed sites of phosphorylation are inhibited by centrinone. Our standard synthetic PLK4 substrate is derived from the activation segment of the CDK activating kinase CDK7, which it phosphorylates on Ser5 in a ‘KSF’ motif, just after the DFG motif (Table 1, Supplementary Fig. 7). This finding raises the possibility that centrinone-mediated inhibition of PLK4 might directly reduce activity of other downstream kinases with site specificities for distinct activation-segment motifs in cells. Potential targets to investigate in the future include CDK7 itself, CDK11, CDK12 and the DNA-damage response modulator CDK18 [91], which all possess centrinone-sensitive phosphorylation sites (Supplementary Fig. 2), and whose substrate specificity (where known) conforms to the Pro-rich motif identified in other proteins found here. However, of the CDK phosphorylation sites that we observed to be modulated by centrinone, none lie within the catalytic (or other known regulatory) domains, and crucially, none are known to regulate protein kinase activity. CHK2 is the only previously defined *in vitro* substrate of PLK4 with protein kinase activity [92] found in our study. However, although it possesses a flexible substrate recognition motif, previous studies have suggested that a basophilic Arg at −3 is preferential for substrate phosphorylation [93], likely ruling out this enzyme as the intermediary of a centrinone-regulated PLK4 network that controls the [pS/pT]P consensus sites reported here. Indeed, the fact that we also identify direct [pS/pT]Pro sites as PLK4 protein targets *in vitro* significantly raises the likelihood that a proportion of the ~300 centrinone inhibited phosphosites identified in cells may well be direct targets of PLK4. The subcellular protein complexes involved in determining these modifications in human cells, and whether they occur co- or post-translationally, remain to be identified.

Multiple members of the CMGC kinase family possess well-documented Pro-directed substrate specificity. Although CDK complexes such as Cyclin B-CDK1 phosphorylate a plethora of crucial M-phase substrates, containing minimal [S/T]P and full consensus [S/T]PXX[L/R] motifs [94–97], it is now clear that at least one CDK enzyme complex can also phosphorylate non-[S/T]P consensus motifs [98], operating through a combined ‘multisite’ code [99], differential substrate co-localization and ordered phosphorylation that is dependent on substrate affinity [100, 101]. Similar findings have also been made with the abscission-controlling PKC-Aurora B complex, whose substrate specificity can switch at different phases of the cell cycle [102]. In the context of PLK4, further studies with full-length proteins are justified to examine whether PLK4 substrate specificity might be altered depending upon the subcellular repertoire of regulatory proteins/substrates formed. Moreover, we speculate that PLK4 substrate-targeting PBDs 1-3 (which are absent in our biochemical analysis, but present in cellular experiments) are worthy of further investigation in order to evaluate differences estabished for PLK4 protein and peptide substrate specificities, either after immunoprecipitation of PLK4 interactomes from cells, or in reconstituted biochemical assays.

## CONCLUSIONS

In this study, we show that exposure of human cells to the PLK4 inhibitor centrinone generates unexpected diverse effects on the phosphoproteome. Our data suggest that the centrinone target PLK4, or a distinct PLK4-regulated kinase(s), phosphorylates [S/T]P motifs on multiple cellular proteins. Notably, protein phosphorylation in this broad [pS/pT]P consensus is markedly downregulated in human cells exposed to centrinone, and PLK4 is able to phosphorylate residues with a Pro at +1 in recombinant proteins (but not derived synthetic peptides) that includes two new PLK4 substrates, PTPN12 and NMYC. In the future, the biological effects of these phosphorylation events will be explored further, in the context of catalytic output of the tyrosine phosphatase PTPN12 and the potential for PLK4 integration into the Aurora A/FBX7/NMYC signaling network. It is anticipated that when assessed alongside complementary PLK4 chemical genetic, depletion or elimination strategies, our proteomic datasets will help define further physiological PLK4 substrates across the cell cycle. These findings will also allow the extent to which a [pS/pT]P phosphorylation consensus represents a direct, indirect or substrate-dependent PLK4 modification in cells, and thus permit the biological roles of PLK4 to be examined in more detail.

## Abbreviations

ATP: adenosine 5’-triphosphate
DMSO: dimethyl sulfoxide
MS: mass spectrometry
MS/MS: tandem mass spectrometry
PBD: polo box domain
PLK: Polo-like kinase
NMYC: N-myc proto-oncogene protein
PTPN12: Protein Tyrosine Phosphatase Non-receptor type 12

---

**Supplementary Figure 1.**
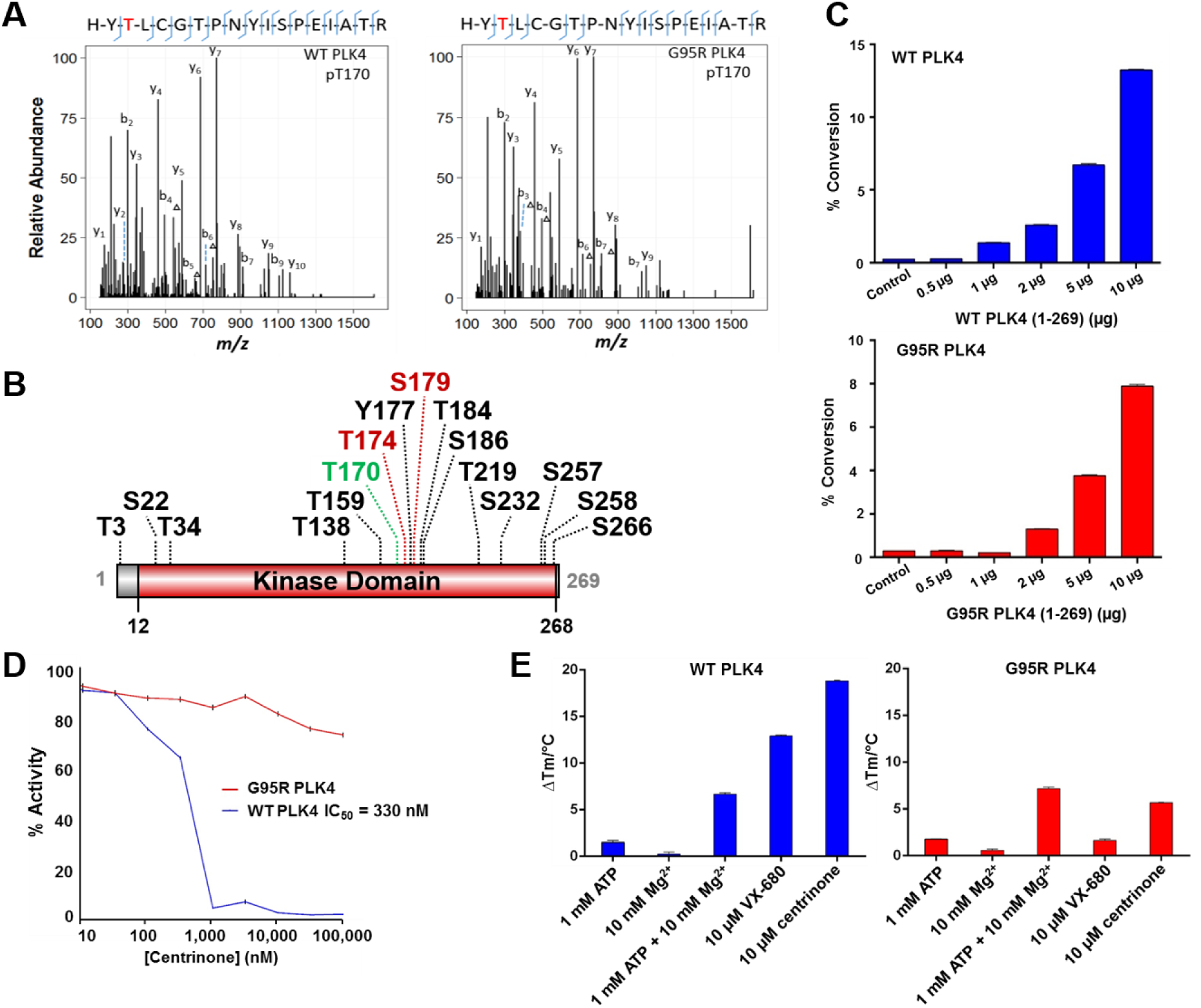
Recombinant WT and G95R PLK4 are catalytically active, autophosphorylating on multiple residues, and G95R PLK4 is highly resistant to inhibition with centrinone. (**A**) Purified recombinant WT or G95R PLK4 (1-269) were digested with trypsin and analysed by LC-MS/MS. MS2 spectra generated by HCD shows expression-induced autophosphorylation at T170 (a known site of phosphorylation required for activity) within the PLK4 activation loop in both recombinant WT & G95R PLK4. (**B**) Autophosphorylation sites identified in WT and G95R PLK4 (1-269) are depicted. T170 is shown in green. Phosphosites in red (T174, S179) conform to a [pS/pT]P consensus. (**C**) Purified recombinant WT or G95R human PLK4 (1–269) were assayed with a fluorescent PLK4 peptide substrate (5’-FAM-FLAKSFGSPNRAYKK) in the presence of 1 mM ATP. (**D**) Purified recombinant WT (blue) or G95R (red) PLK4 (1-269) were incubated with fluorescent peptide substrate in the presence of DMSO (control) or the indicated concentrations of centrinone and 1 mM ATP. The extent of peptide phosphorylation (converted to activity) was analysed by mobility shift assay using the EZ Reader platform. Data in (**C**) are from triplicate assays performed twice. Data in (**D**) are from a single triplicate assay. Similar results were seen in a separate experiment. (**E**) DSF analysis of WT (blue) or G95R (red) PLK4 in the presence of the indicated concentrations of nucleotides, metals or inhibitor compounds.

**Supplementary Figure 2.**
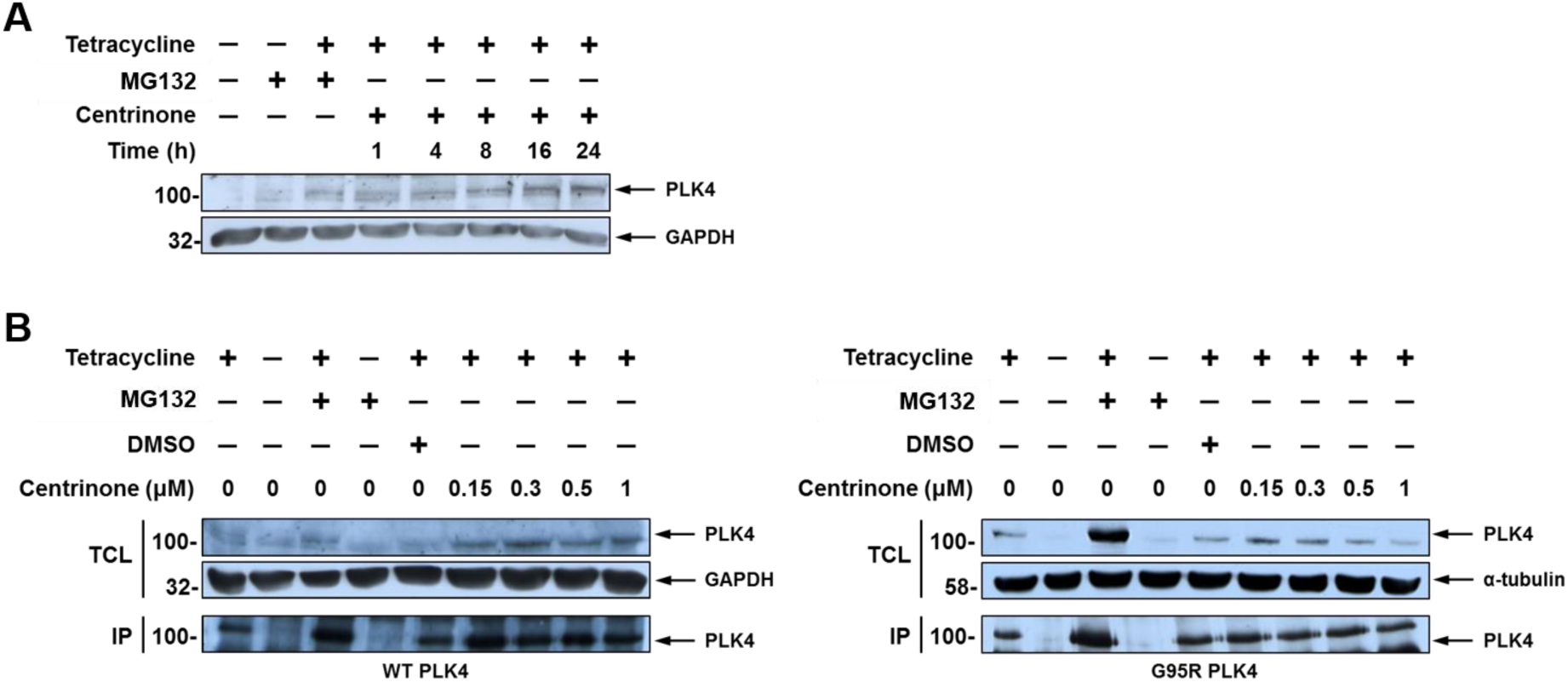
FLAG-WT PLK4 but not FLAG-G95R PLK4 is stabilised by centrinone in a concentration-dependent manner. Expression of FLAG-WT PLK4 or FLAG-G95R PLK4 was induced with 1 μg/mL tetracycline for 18 hours. Cells were incubated with 10 μM MG-132 for 4 hours and either (**A**) 150 nM centrinone for the times indicated, or (**B**) with the indicated concentrations of centrinone for 4 hours. Total cell lysates (TCL) and immunoprecipitated FLAG-PLK4 were analysed by western blotting using the indicated antibodies.

**Supplementary Figure 3.**
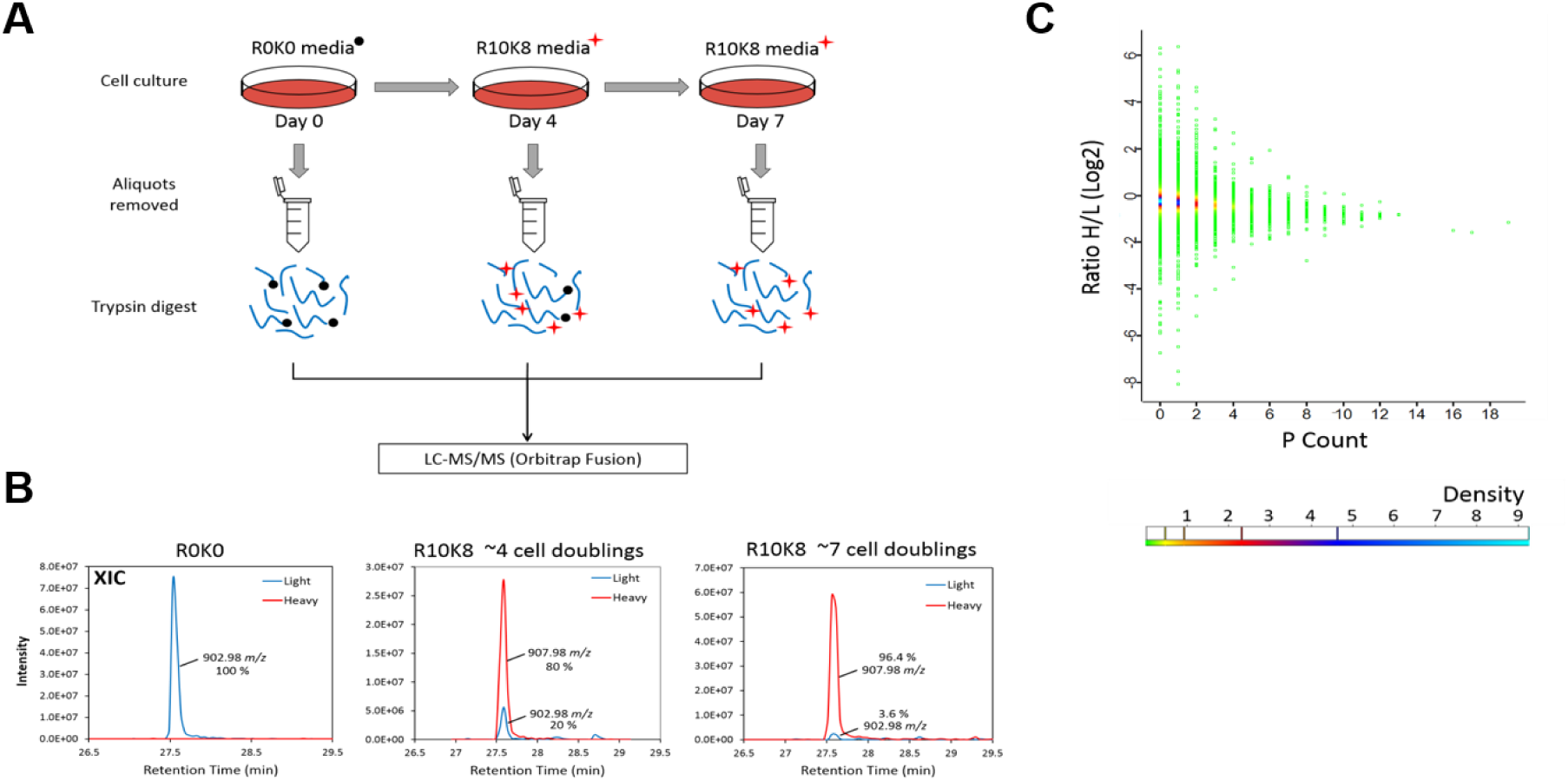
Efficient SILAC labelling of U2OS-FLAG WT PLK4 cells. (**A**) U2OS cells cultured in ‘light’ media were sub-cultured into ‘heavy’ media (R10K8) containing the stable isotopes arginine (^13^C_6_ ^15^N_4_) and lysine (^13^C_6_ ^15^N_2_), allowing incorporation of the isotope-labelled amino acids into newly synthesised proteins during cell growth and protein turnover. At each passage, an aliquot of cells were removed for analysis by LC-MS/MS following tryptic proteolysis, to assess incorporation of the heavy labelled amino acids. (**B**) The extracted ion chromatograms (XIC) show the ion signals for an exemplar doubly charged unlabelled tryptic peptide ion at m/z 902.98 unlabelled. (Left) XIC of the light (unlabelled) peptide in blue, with increasing amount of ‘heavy’ labelled peptide ion (m/z 907.98; 10 Da mass difference) being observed after ~4 and then 7 cell doublings (red), at which point labelling has reached over 96%. (**C**) A density plot was generated to assess the metabolic conversion of Arg to Pro. Non-normalised H/L ratios were plotted against the total proline count from all identified peptides. The data points are colour coded based on density. No global drift toward the unlabelled peptides were observed, confirming no significant metabolic conversion.

**Supplementary Figure 4.**
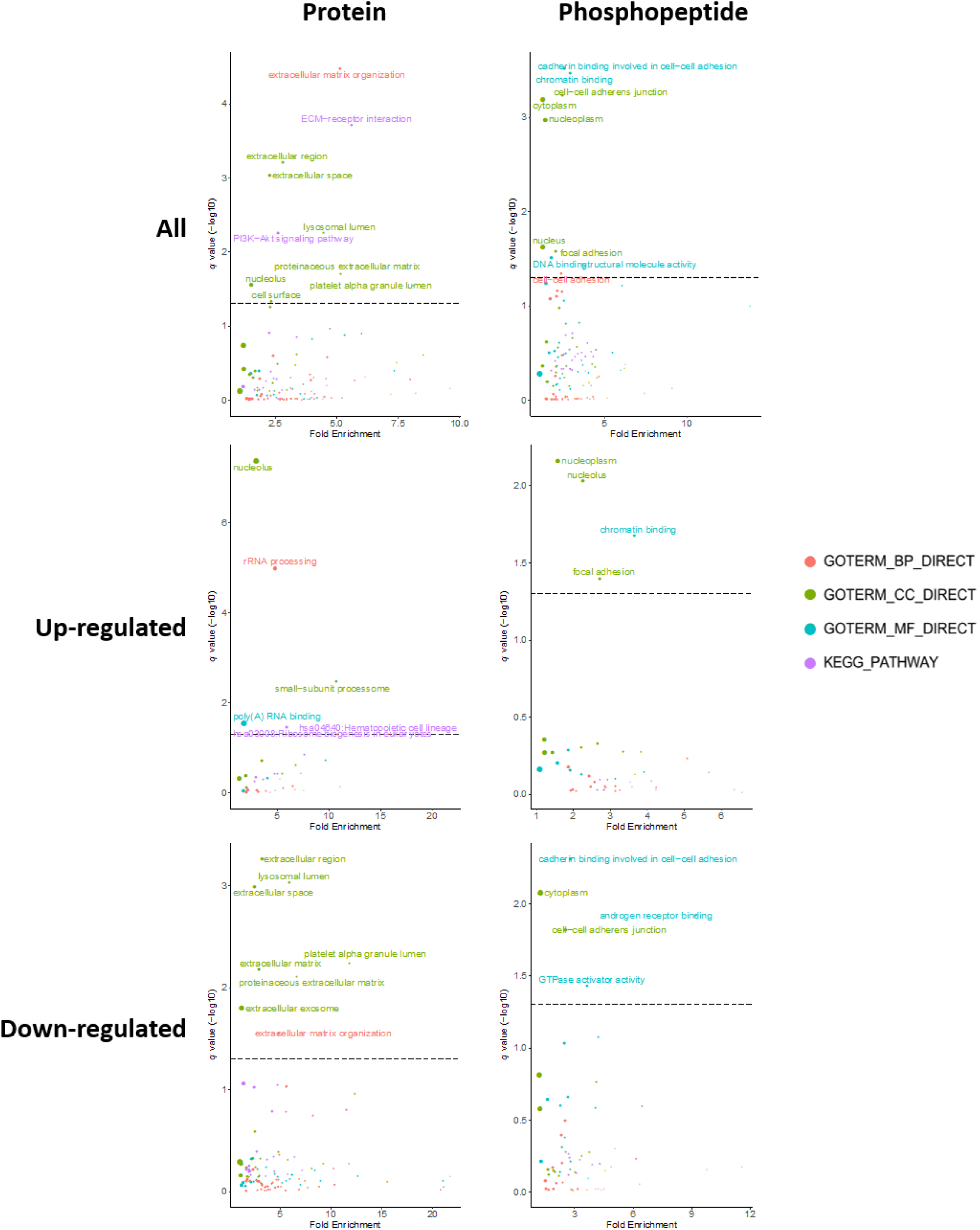
GO term enrichment analysis. Data for significantly regulated protein expression (top), or proteins with differentially regulated phosphopeptides (bottom) observed after centrinone treatment of FLAG-WT PLK4 U2OS cells. Proteins/phosphopeptides with a Benjamini-Hochberg adjusted p-value ≤0.05 are labelled. BP = biological process (red); CC = cellular compartment (green); MF = molecular function (cyan); KEGG pathway (purple). The size of the node is representative of the number of proteins contributing to a select category.

**Supplementary Figure 5.**
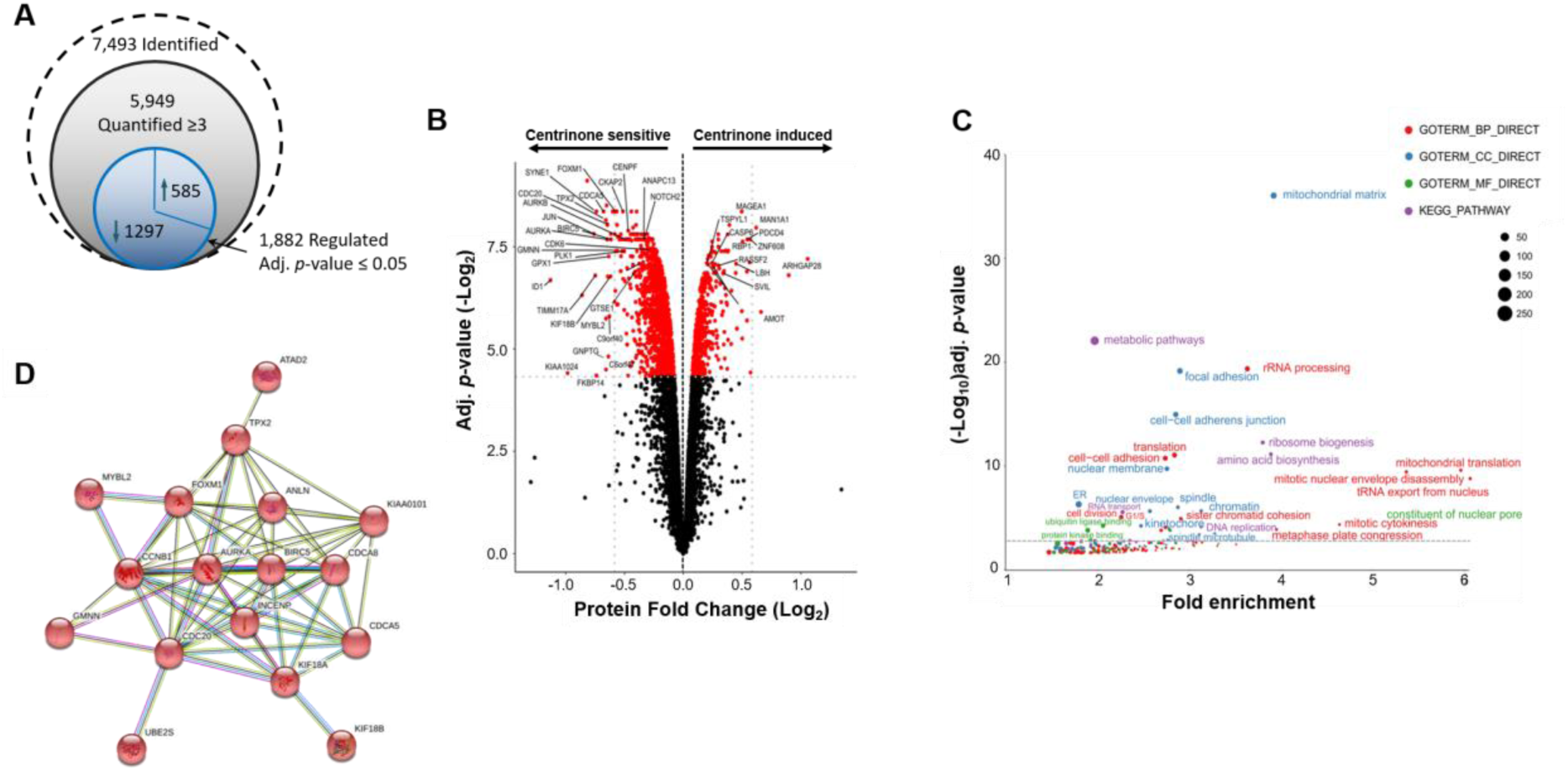
Significant protein regulation is observed in FLAG-G95R PLK4 cells following centrinone treatment. (**A**) Total numbers of identified, quantified and differentially regulated proteins are indicated. (**B**) Volcano plot showing protein fold changes following Bayesian statistical analysis to evaluate significant differences. Log_2_-fold change (Heavy/Light) are presented as a function of the –Log_2_ Benjamini-Hochberg adjusted p-value; those proteins with an adjusted p-value ≤0.05 are highlighted in red. Select data points are annotated with their protein accession number. (**C**) GO term enrichment analysis of significantly regulated proteins using DAVID. Proteins/phosphopeptides with a Benjamini-Hochberg adjusted p-value ≤0.05 are labelled. BP = biological process (red); CC = cellular compartment (blue); MF = molecular function (green); KEGG pathways (purple). (**D**) STRING interaction analysis reveals a network of down-regulated mitotic proteins.

**Supplementary Figure 6.**
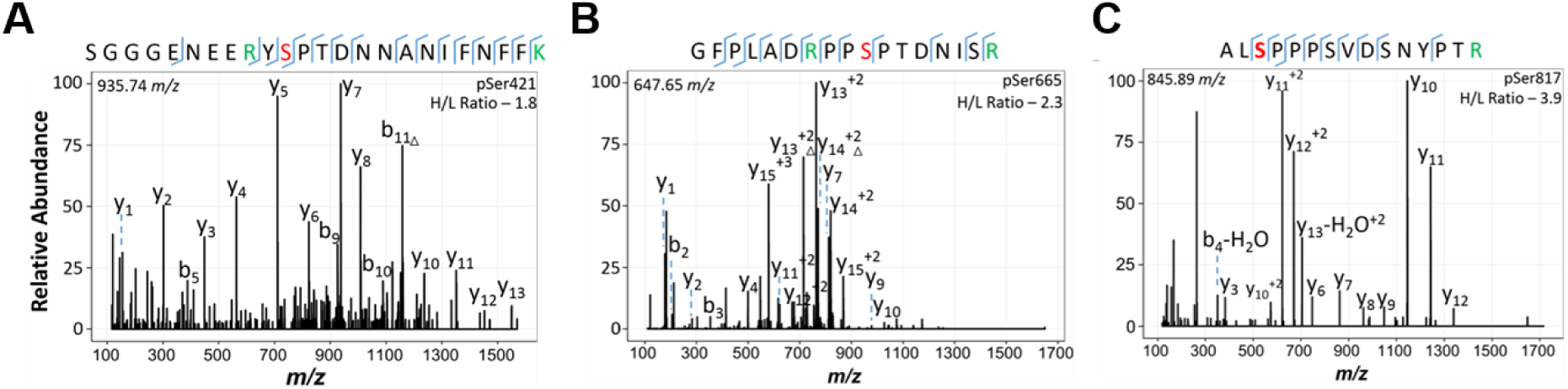
Centrinone-mediate upregulation of PLK4 phosphosites. Three significantly up-regulated (adj. p value ≤0.05) PLK4 phosphosites were identified from U2OS FLAG-WT PLK4 cells following treatment with centrinone (300 nM centrinone for 4 hours). The peptide sequence and the identified phosphosite (red) is detailed on each tandem mass spectrum. SILAC labelled K/R residues are in green. (**A**) Doubly charged ion at m/z 935.74, identifying pSer421 as the site. (**B**) Doubly charged ion at m/z 647.65, identifying pSer665. (**C**) Doubly charged ion at m/x 845.89, identifying pSer817 as the site.

**Supplementary Figure 7.**
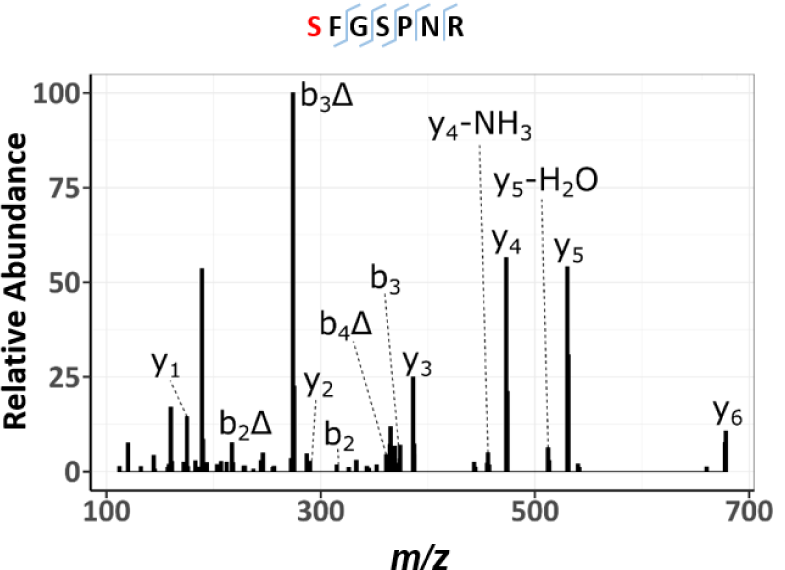
HCD product ion mass spectrum of the digested FLAKSFGSPNRAYKK peptide substrate. Fluorescently-labelled FLAKSFGSPNRAYKK peptide was incubated with PLK4 in 25 mM HEPES (pH 7.4), 1 mM ATP, 5 mM MgCl_2_, and 0.001% (v/v) Brij 35 at 37°C. Reactions were then digested with trypsin to remove the fluorescent tag, and the digest subjected to LC-MS/MS analysis. Note the digested peptide yields the sequence SFGSPNR with a precursor m/z value of 422.6711. Red denotes the site of phosphorylation. Δ indicates H_3_PO_4_ loss.

## ACKNOWLEDGEMENTS

We thank Dr Gopal Sapkota (University of Dundee) for the generous donation of parental U2OS cells. The authors also wish to thank Prof. Sonia Rocha for help with FACS analysis, Sam Evans for media preparation, Alex Holme for technical support, and the Centre for Cell Imaging at the University of Liverpool.

## AUTHOR CONTRIBUTIONS

DPB, PAE and CEE obtained funding, designed the experiments and analysed the data alongside CC and AC. AK, SP and ARJ performed bioinformatics analysis. CC, AC and PJB performed MS analysis, and DM and staff in the Centre for Cell Imaging helped with immunofluorescence studies. PAE and CEE wrote the paper, with contributions from all authors, who also approved the final version prior to submission.

## DATA AND MATERIALS AVAILABILITY

All data needed to evaluate the conclusions made are available in the main or supplementary sections of the paper. The mass spectrometry proteomics data have been deposited to the ProteomeXchange Consortium via the PRIDE partner repository [103] with the dataset identifier PXD018704 and 10.6019/PXD018704.

## SUPPLEMENTARY FIGURE LEGENDS

**Supplementary Table 1. Proteome changes induced by centrinone.**

List of the identified proteins in the FLAG-WT PLK4 (sheet 1) or the FLAG-G95R PLK4 (sheet 2) cell line and the fold change (FC) in response to centrinone treatment. Statistical analysis was performed using the LIMMA package in R. *p-*value and adjusted p-values calculated using the Benjamini-Hochberg correction for multiple testing) are detailed for each protein family.

**Supplementary Table 2. Phosphoproteome changes induced by centrinone.**

List of the identified phosphopeptides, the sites within the proteins and the PTM-score computed site localisation confidence in the FLAG-WT PLK4 (sheet 1) or the FLAG-G95R PLK4 (sheet 2) cell line, and the fold change (FC) in response to centrinone treatment. Statistical analysis was performed using the LIMMA package in R. *p-*value and adjusted p-values calculated using the Benjamini-Hochberg correction for multiple testing) are detailed for each protein family. Also detailed are the site conservation across 100 species (listed in sheet 3) and the prevlance of prior observation in either PhosphositePlus (PSP) or Peptide Atlas (PA) – see Methods for detailed information.

